# Poincaré Maps for Analyzing Complex Hierarchies in Single-Cell Data

**DOI:** 10.1101/689547

**Authors:** Anna Klimovskaia, David Lopez-Paz, Léon Bottou, Maximilian Nickel

## Abstract

The need to understand cell developmental processes spawned a plethora of computational methods for discovering hierarchies from scRNAseq data. However, existing techniques are based on Euclidean geometry, a suboptimal choice for modeling complex cell trajectories with multiple branches. To overcome this fundamental representation issue we propose Poincaré maps, a method that harness the power of hyperbolic geometry into the realm of single-cell data analysis. Often understood as a continuous extension of trees, hyperbolic geometry enables the embedding of complex hierarchical data in only two dimensions while preserving the pairwise distances between points in the hierarchy. This enables direct exploratory analysis and the use of our embeddings in a wide variety of downstream data analysis tasks, such as visualization, clustering, lineage detection and pseudo-time inference. When compared to existing methods —unable to address all these important tasks using a single embedding— Poincaré maps produce state-of-the-art two-dimensional representations of cell trajectories on multiple scRNAseq datasets. More specifically, we demonstrate that Poincaré maps allow in a straightforward manner to formulate new hypotheses about biological processes unbeknown to prior methods.

**Significance statement:** The discovery of hierarchies in biological processes is central to developmental biology. We propose Poincaré maps, a new method based on hyperbolic geometry to discover continuous hierarchies from pairwise similarities. We demonstrate the efficacy of our method on multiple single-cell datasets on tasks such as visualization, clustering, lineage identification, and pseudo-time inference.

## Introduction

Understanding cellular differentiation, e.g., the transition of immature cells into specialized types, is a central task in modern developmental biology. Recent advances in single-cell technologies, such as single-cell RNA-sequencing and mass cytometry, enabled important insights into these processes based on high-throughput cell measurements^1–4^. Computational methods to accurately discover and represent cell development processes from large datasets and noisy measurements are therefore in great demand. This is a challenging task, since methods are required to reveal the progression of cells along continuous trajectories with tree-like structures and multiple branches (e.g., as in Waddington’s classic epigenetic landscape^5^). Multiple advances have been made towards this goal of discovering and analyzing hierarchical structures from single-cell measurements^6^. In particular, methods leveraging assumptions about hierarchical structure for visualization^7–12^, clustering^13;14^, and pseudo-time inference^15;16^, have fueled unprecedented successes in developmental biology. To visualize hierarchical relationships in cell development, many state-of-the-art methods embed cell measurements in low-dimensional Euclidean spaces^7;8;17;18^. However, this approach is limited when modeling complex hierarchies, as low-dimensional Euclidean embeddings distort pairwise distances between measurements substantially. The resulting embeddings are problematic not only for visualization, but also for other downstream tasks such as clustering and lineage identification.

To overcome the issues of data dimensionality reduction in Euclidean spaces, we propose Poincaré maps, a novel method to compute embeddings in hyperbolic spaces. These enable multiple advantages. First, hyperbolic spaces can be thought of as a continuous analogue to trees and enables low-distortion embeddings of hierarchical structures in as few as two dimensions^19^. Second, the metric structure of hyperbolic spaces retains the ability to model continuous trajectories using pairwise distances of measurements, and allows to employ the obtained embeddings in downstream tasks such as clustering, lineage detection and pseudo-time inference. Third, the Riemannian structure of hyperbolic manifolds enables the use of gradient-based optimization methods what is essential to compute embeddings of large-scale measurements. Fourth, while we follow Nickel et al.^20^ to leverage the Poincaré disk as an embedding space, we are first to employ pairwise distances obtained from a nearest-neighbor graph as a learning signal to construct hyperbolic embeddings for the discovery of complex hierarchies in data.

An important property of Poincaré maps is that it allows to approach all these different tasks using a *single embedding*, by combining the identification of clusters, trajectories, and hierarchies in an unsupervised manner. To the best of our knowledge, this is not possible with existing methods, which we review in the following. t-SNE^17^ is a state-of-the-art visualization method that exploits local similarities to achieve visual separation of the clusters in the data. However, t-SNE does not preserve global similarities between clusters, and therefore does not guarantee that the global hierarchical structure will be preserved. UMAP^18^ computes a low-dimensional Euclidean representation of data that preserves the topological structure. However, there are no guarantees that there exists a low-dimensional representation of complex tree topologies in a two-dimensional Euclidean space. Diffusion maps^7^ specifically tackles the problem of capturing diffusion-like dynamics and continuous branching in the data. However, it allows to visualize only simple branching structures in two dimensions. Graph abstractions^21^ (PAGA) and Monocle 2^15^ are another class of methods to capture and visualize hierarchical relationships in the data. PAGA produces an “abstracted graph” with nodes corresponding to partitions of the data, and edges representing relationships between these nodes. PAGA does not represent the relationships within partitions. However, PAGA can be used to initialize UMAP and ForceAtlas2, as done by the authors. Despite the fact that ForceAtlas2^22^ produces a good visual layout of a tree topology, it does not preserve hierarchical distances. PHATE^23^, a method that has been demonstrated able to recover hierarchies with multiple branches, is also affected by the distortion artifacts of Euclidean spaces. Monocle 2^15^ forces a tree-like topology on the data using “reversed graph embedding” in a low-dimensional Euclidean space. However, *similar to* UMAP, such a representation might not exist for complex trees. SIMLR^10^ is a multi-kernel learning designed to perform well on datasets with multiple clusters, making it a poor choice to model data with continuous trajectories. Finally, SAUCIE^10^ is an autoencoder model, which is optimized through reconstruction error, therefore its properties for preserving local and global similarities are theoretically less understood.

## Results

### Preservation of high-dimensional hierarchies through Relative Forest Accessibilities index in a Poincaré disk

Our method, *Poincaré maps*, is inspired by ideas from manifold learning and pseudo-temporal ordering^24;25^. Given feature representations of cells such as their gene expressions, we aim to estimate the structure of the underlying tree-like manifold in three main steps (**Fig. 1** and Methods): First, we compute a *connected k*-nearest neighbor graph (*k*NNG)^26^ where each node corresponds to an individual cell and each edge has a weight proportional to the Euclidean distance between the features of the two connected cells.1 To enforce connectivity we propose a simple procedure, described in Online Methods. The purpose of this first step is to estimate the local geometries of the underlying manifold, around which Euclidean distances remain a good approximation. Second, we compute global geodesic distances from the *k*NN graph, by traveling between all pairs of points along the weighted edges. This step can be computed efficiently using all pairs of shortest paths, or related measures such as the “Relative Forest Accessibilities” (RFA) index^27^. The purpose of this second step is to estimate the intrinsic geometry of the underlying manifold. These two first steps are commonly used in manifold learning to approximate the structure of an unknown manifold from similarities in the feature space^16;26;28;29^. As a third step, we compute a two-dimensional embedding per cell in the Poincaré disk, such that their hyperbolic distances reflect the inferred geodesic distances. The geometry of the Poincaré disk allows us to model continuous hierarchies efficiently. More specifically, embeddings that are close to the origin of the disk have a relatively small distance to all other points, representing the root of the hierarchy, or the beginning of a developmental process. On the other hand, embeddings that are close to the boundary of the disk, have a relatively large distance to all other points and are well-suited to represent leaf nodes. Thus, in Poincaré embeddings, we expect that nodes with small distances to many other nodes will be placed close to the origin of the disk. While such cells are likely from an early developmental stage, they do not necessarily belong to the root of the hierarchy (**Supplementary Fig. 4–6**). When a cell belonging to the root is known, we perform a translation on the Poincaré disk to place this cell in the center of the disk, easing the visualization of the hierarchy (see **Methods**).

**Figure 1.**
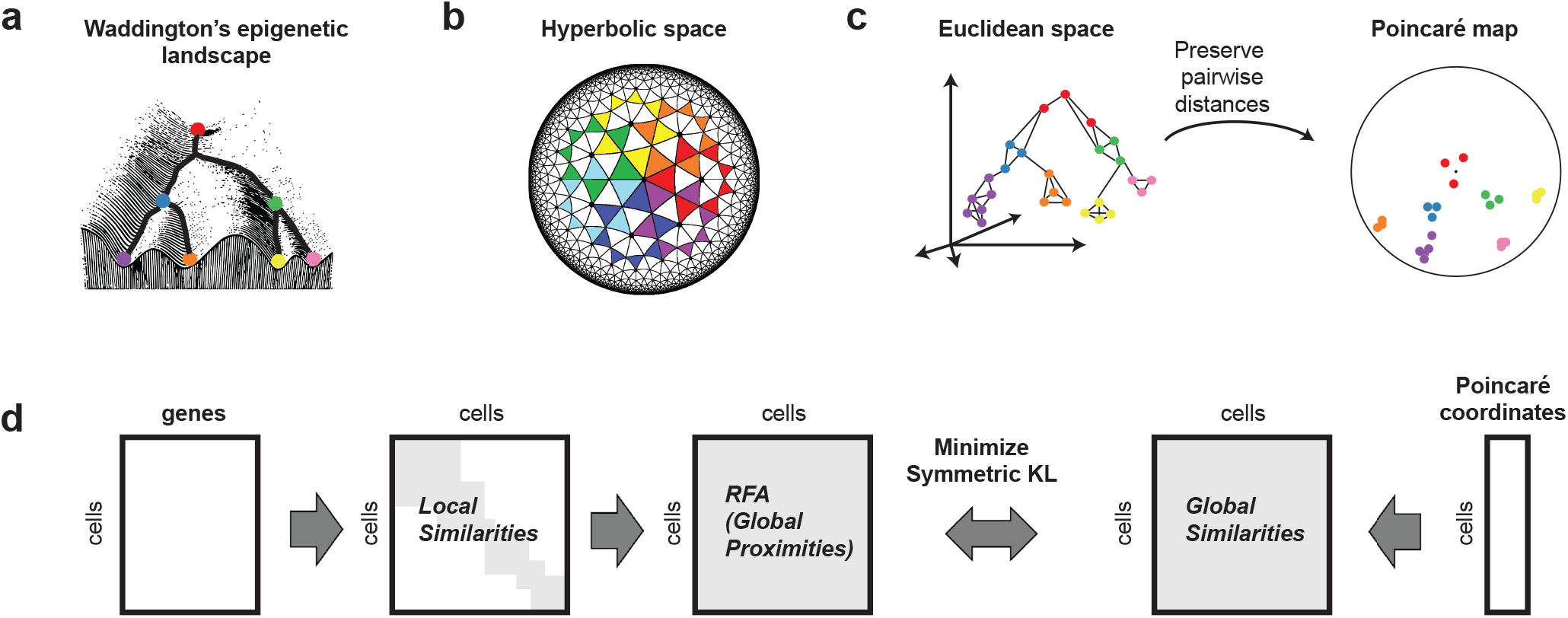
*Poincaré maps* discover hierarchies and branching processes. (**a**) Our goal is to recover cell developmental processes, depicted here on the Waddington landscape. (**b**) Poincaré disks provide a natural geometry to preserve hierarchical structures and pairwise similarities in two dimensions. Poincaré disks grow as we approach their boundary: all the triangles depicted in the figure are of equal size. (**c**) Poincaré maps first estimate geodesic distances, computed from a connected k-nearest neighbor graph. Second, they compute two-dimensional hyperbolic embeddings that preserve these similarities. (**d**) Overview of *Poincaré maps* embedding procedure. From a given feature matrix, *Poincaré maps* firsts estimates local similarities based on a user specified local distance metric (Euclidean, cosine, etc.) and Gaussian kernel with a tunable parameter *σ*. Local similarities then used to compute global proximities on the dataset. By means of Riemanninan optimization of KL divergence, global proximities are aimed to be preserved through global distances in Poincaré disk.

Poincaré maps have several hyper-parameters to tune, such as the number of nearest neighbors (*k*), the bandwidth of the local kernel to convert distances into similarities (*σ*), and the scaling parameter to compute similarities in the Poincaré disk (*γ*). **Supplementary Fig. 2** demonstrates the performance of Poincaré maps with respect of the choice of these hyper-parameters and across different random seeds.

### All-in-one: visualization, clustering, lineage detection and pseudo-time inference

In the following, we compare Poincaré maps to prior state-of-the-art methods on various single-cell analysis tasks: visualization and lineage detection (PCA, Monocle 2^15^, PAGA^21^, diffusion maps^7^, t-SNE^17^, UMAP^18^, ForceAtlas2^22^, SAUCIE^12^, PHATE^23^ and SIMLR^10^), clustering (Louvain^30^, agglomerative, k-means) and pseudo-time inference (diffusion pseudo-time^16^).

For this purpose, we employ multiple synthetic datasets generated from known dynamical systems and four single-cell RNA sequencing datasets varying in size, complexity (number of cell types and branches) and single-cell technology to acquire the data^2;3;31;32^. We compare Poincaré maps with the canonical hematopoetic cell lineage tree^33^, and various state-of-the-art embeddings (**Supplementary Note 2**).

First, we evaluate the capabilities of Poincaré maps for data visualization and dimensionality reduction. It is not possible for humans to comprehend visualizations in more than three dimensions, and a third dimension already adds additional challenges for interpretation. However, for existing methods, even three-dimensional embeddings are not sufficient to capture the underlying manifold structure on many complex hierarchies. Here, we demonstrate that as few as two dimensions of Poincaré maps are enough to reconstruct the hierarchy and preserve global similarities on the datasets with a very high complexity. A common way to evaluate the quality of an embedding in labeled datasets is to use classification scores. However, this evaluation approach has limitations in the context of single-cell data, and specifically for recovering hierarchies and continuous developmental trajectories. First, quite often labels are assigned using some unsupervised learning approach, such as clustering. This could promote the dimensionality reduction method that better agrees with the label assignment method, rather than with the objective ground-truth. Second, discrete labels do not easily apply to datasets with continuous trajectories, where a clear-cut cutoff between cell types does not exist. Third, it contains no information about the quality of preservation of global similarities, e.g. positions of clusters relative to each other in the hierarchy. Instead, we use a scale-independent quality criteria^34^ (see Methods and **Supplementary Note 2**), which was demonstrated to be a good metric to compare embeddings in an unbiased way. The criteria consists of estimating two scalar values *Q*_*local*_ and *Q*_*global*_ reflecting local and global properties of the dataset. We follow the assumption of Lee et al.^34^ that a single-cell dataset comprises a smooth manifold and a good dimensionality reduction method would preserve local and global distances on this manifold.

An important result from our experiments is that Poincaré maps was the only method that demonstrated the ability to visualize the correct branching structure of developmental processes for *all* datasets in terms of this quality metric (**Fig. 3**, **Supplementary Fig. 2**). Separate visual comparison of various embeddings (**Supplementary Fig. 4 – 10**) demonstrates the superior readability advantages of Poincaré maps. For example, on the dataset Paul et al.^2^ only Poincaré maps and t-SNE identify the lymphoid cluster, while this important population remained invisible during exploratory data analysis when using UMAP or ForceAtlas2. Although t-SNE visualizes separate clusters well for Paul et al. dataset, it disregards the hierarchical structure between clusters (see also example in **Supplementary Fig. 8**). Knowledge of the position of a newly identified cluster in the developmental hierarchy could be further exploited for assigning labels (e.g. “lymphoid population”) or, when the population was not known, for designing experiments to test morphological properties. Finally, Poincaré maps places the 16Neu cluster downstream of 15Mo in the hierarchy —in contrast to the canonical hierarchy, where neutrophils and monocytes are located at the same level. This result is in line with the analysis of Wolf et al.^21^, indicating that the inconsistency is due to a faulty labeling of the clusters.

A back-to-back comparison of a quality metric with visual inspection gives a good intuition about the meaning of the metric quality scores and embedding properties. In particular, on datasets with simple trajectories (e.g., ToggleSwitch, MyeloidProgenitors), methods such as PCA or SAUCIE show strong performance since they preserve local similarities well. However, the performance of these methods drops significantly for the datasets with many different cell types and branches, such as Planaria and C. elegans. Poincaré maps in their turn, don’t suffer from this limitation (**Fig. 3**) and significantly outperforms other methods in terms of both local and global metrics. This allows us to *summarize the whole C. Elegans cell atlas in a single Poincaré maps embedding* (**Fig. 2**). This was not possible with UMAP with any choice of parameters, as reported by the authors in their original study. This makes Poincaré maps a strong candidate for visualization of single-cell atlases. Additional analysis of the age of the embryo on Poincaré maps revealed two distinct populations of germlines. One of these subpopulations is placed close to the border of the disc and more close to mature cell types, which potentially reflects transcription diversity of this subpopulation from other cells at early stages. The second subpopulation is close to other cells at the early stage. We randomly picked up a cell from the second subpopulation, and assigned it at as a root. **Fig. 2 (c)** demonstrates the relative positioning of the cell types in the hierarchy and comparison of the Poincaré pseudo-time to the age of the embryo. We can see that it agrees with the age of embryo quite well, except for very early stages (<130). However, lineages are not perfectly synchronized, therefore we see significant variability on the plot.

**Figure 2.**
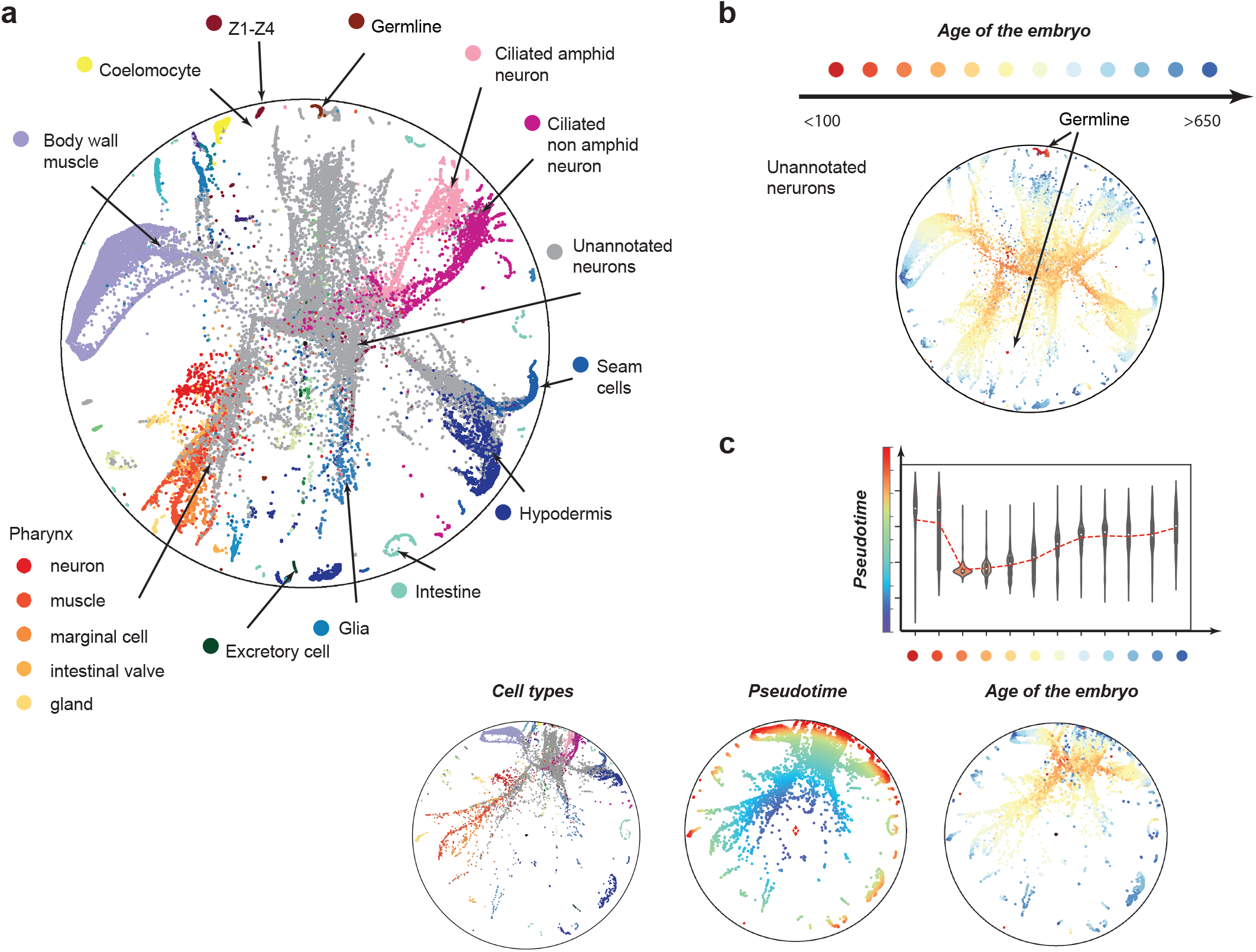
Analysis of C. elegans cell atlas. (**a**) Poincaré map (without rotation) on a 40,000 cell random subsample and 100 PCA components. The parameters used for embedding are (*k* = 15, *σ* = 2.0, *γ* = 3.0). Main cell types are annotated with a text legend, the rest are separated by color. (**b**) Poincaré maps places mature cell types towards the border of the disk. Two subpopulations of germline cells are apparent from the embedding. (**c**) Rotation and comparison of Poincaré maps with respect to randomly picked up root cell form one of the sub-populations of the germlines (the one that is more similar to the rest of cell types of early the age the embryo). Red line is an average pseudo-time distance for a given age of the embryo.

**Figure 3.**
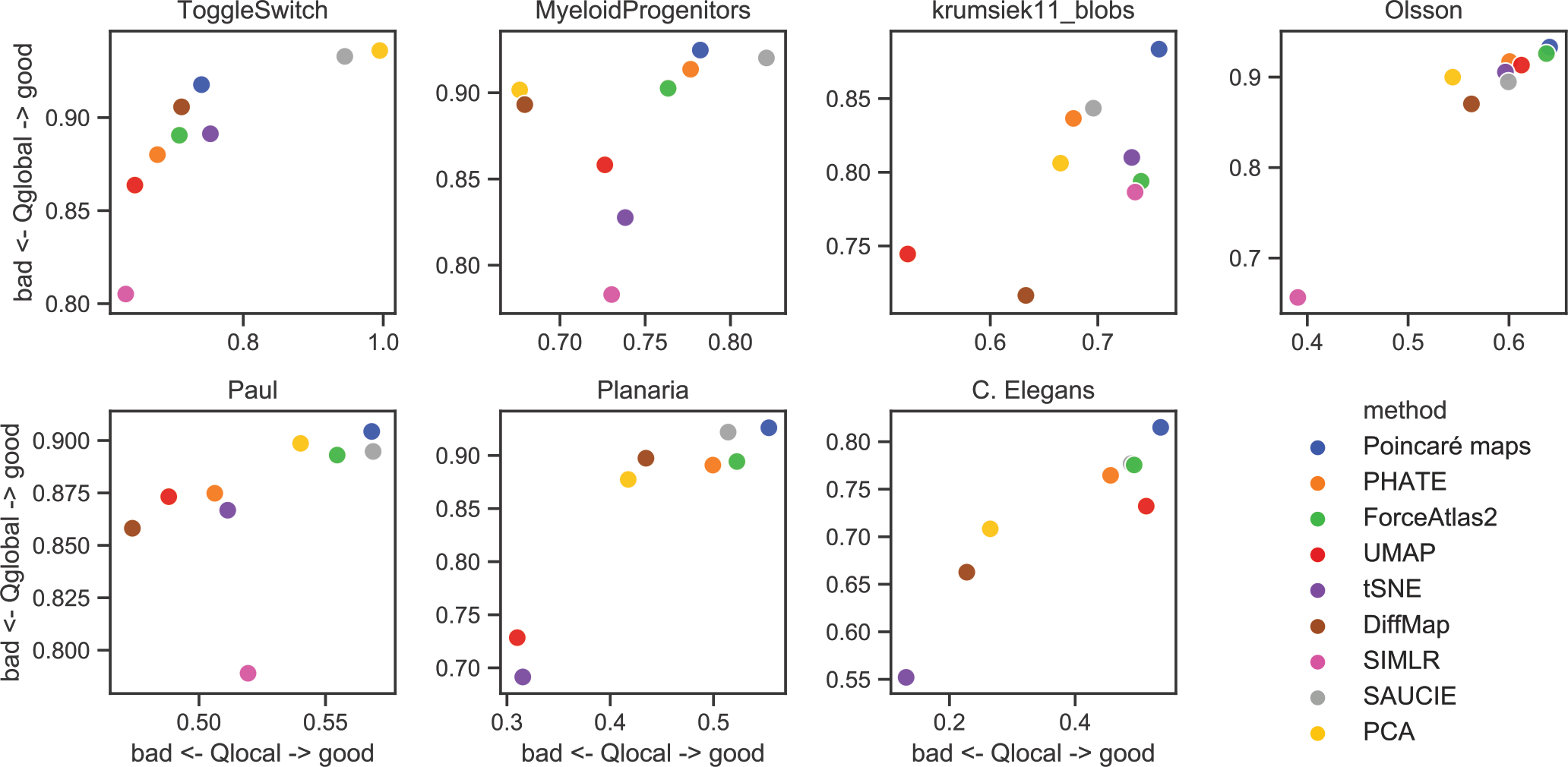
Comparison of embedding quality metric (best case) for various datasets. Poincaré maps perform consistently well on all synthetic and real-world datasets in our evaluation. Planaria and C. elegans datasets —which exhibit the highest complexity in terms of number of branching trajectories— are datasets where Poincaré maps perform significantly better than other methods.

In addition to these results, we demonstrate in **Supplementary tables 1–2** that Poincaré maps could be directly applied to achieve state-of-the art results on clustering and pseudo-time inference. Notably, for pseudo-time inference the results are comparable with diffusion pseudo-time, but in Poincaré maps these clusters are directly accessible as a distances from root node. Therefore they are not only fast to compute given the embedding, but also allow to intuitively interpret a Poincaré maps plot with the root node in the center of the Poincaré disk.

### Poincaré maps generate new hypothesis about early blood development in mice

As a deeper case study, we analyze the dataset of early blood development in mice, previously studied by Moignard et al.^1^, using Poincaré maps. This dataset contains measurements of cells captured *in vivo* with qRT-PCR at different development stages: primitive streak (PS), neural plate (NP), head fold (HF), four somite GFP (Runx1) negative (4SG−) and four somite GFP positive (4SG+) (**Fig. 4 (a)**). The stages correspond to different physical times of the experiment between embryonic day 7 and day 8.25. We compare our results obtained with Poincaré maps to Moignard’s diffusion maps study^1^, and to Haghverdi’s reconstruction of diffusion pseudo-time^16^. Poincaré maps provide a qualitatively different visualization of the developmental process, where we are able to visualize the whole spectrum of the heterogeneity arising from the onset of the process. Neither PCA nor diffusion maps are able to provide a visualization of this process. While Moignard’s and Haghverdi’s analyses suspected an asynchrony in the developmental process, neither their application of PCA or diffusion maps were able to reveal this. In particular, previous studies suggest that the split into endothelial and erythroid sub-populations happens in the head fold. Our analysis using Poincaré maps indicates that the sub-population fate of the cells is already predefined at primitive strike. Additionally, Poincaré maps reveal a separate cluster consisting of a mixture of cells at different developmental stages (**Supplementary Fig. 12**). This cluster is referred to as “mesodermal” cells by Moignard et al., while by Haghverdi et al. considers it as the root of the developmental process. However, as we demonstrate in **Supplementary Fig. 13 – 14**, assigning this cluster as the root of the hierarchy would lead to a contradiction with the physical direction of time. By virtue of the Poincaré visualization, we reassigned the root of the developmental process to the furthest PS cell not belonging to the “mesodermal” cluster. We picked up a root cell from PS as to ease clustering by angle for lineage detection. More specifically, we chose the most “exterior” cell from the PS cluster, by visual inspection. Given our reassigned root, we separate the dataset into five potential lineages (see **Methods**), to find the asynchrony in the developmental process in terms of marker expressions (**Fig. 4 (b)**). Analysis of the composition of cells belonging to each lineage (**Fig. 4 (c)**) indicates that erythroid cells belong only to lineage 0 and this lineage contains no endothelial cells. **Fig. 4 (d)** shows a substantially improved agreement of Poincaré pseudo-time (with the newly reassigned root) with the experimental time (stages) compared to the pseudo-time ordering proposed by Haghverdi et al. The analysis of gene expressions of main endothelial and hemogenic markers agrees with the known pattern of gene activation for endothelial and erythroid branches (**Supplementary Fig. 14 – 15**). **Fig. 4 (e)** also demonstrates that the main hemogenic genes for the erythroid population are already expressed at the PS stage (details in **Supplementary Note 3**) and that the differences in gene expression apparent at all the stages between the lineages. Our analysis using Poincaré maps suggests therefore that the fate of erythroid and endothelial cells could already be defined at primitive streak.

**Figure 4.**
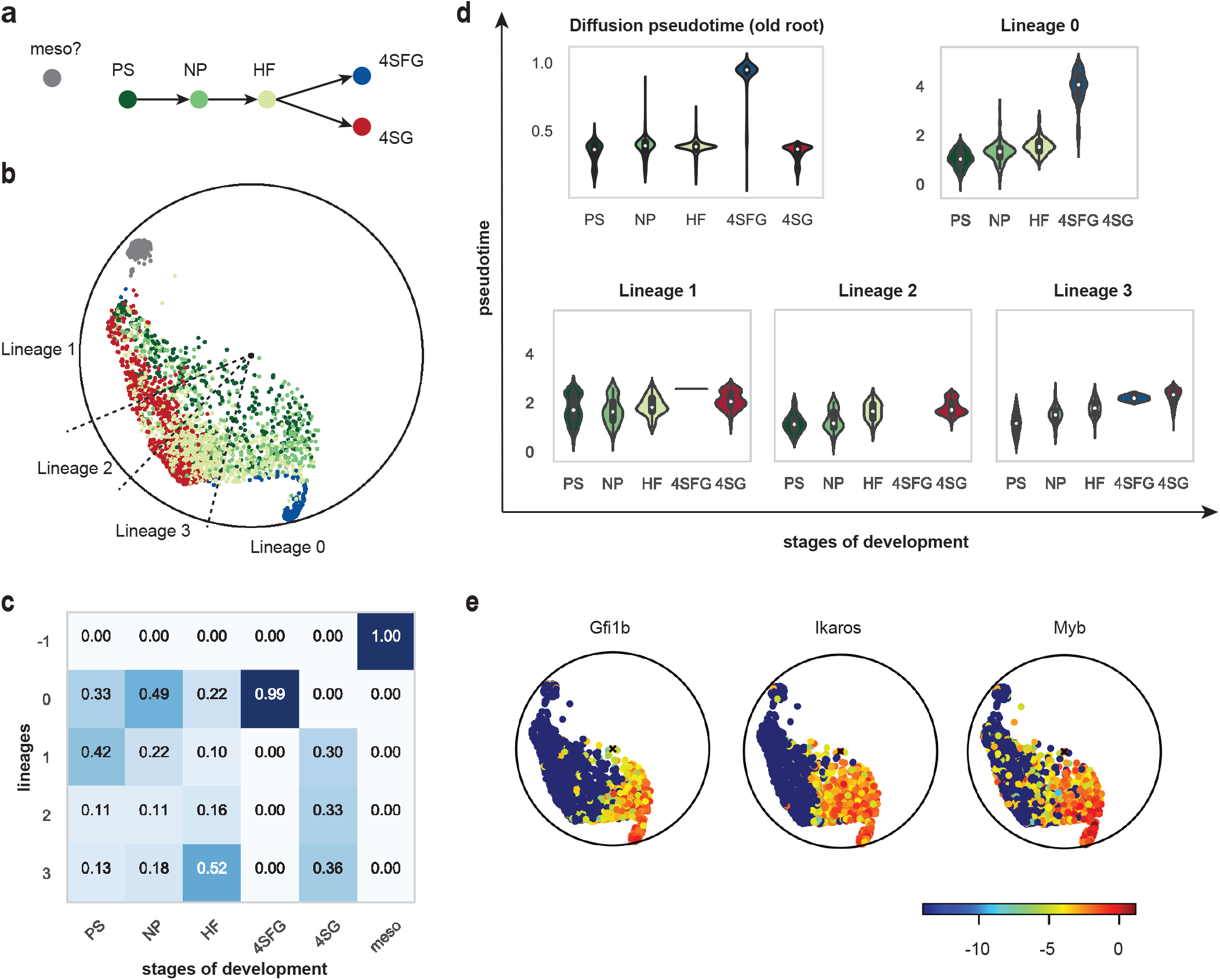
(**a**) Developmental hierarchy proposed by Moignard et al. and Haghverdi et al. (**b**) Rotated Poincaré map with respect to reassigned root. Grey cluster represent a cluster of potential outliers or “mesodermal” cells as suggested by Moignard et al. Lineage slices were obtained with Poincaré maps (see Methods). (**c**) Composition of detected lineages in terms of the presence of cells from different developmental stages. (**d**) The ordering of cells proposed by Poincaré maps here has a much better agreement with developmental stages than ordering originally proposed by Haghverdi et al.: we see a very clear correlation of Poincaré pseudo-time with actual developmental time. (**e**) Gene expression of main hemogenic genes. Hemogenic genes of erythroid lineage are already expressed at the PS and NP stages.

## Discussion

The rapid onset of popularity and accessibility of single-cell RNA sequencing technologies fueled the development of new computational approaches to analyze these data. While many computational methods exist, their results often disagree between each other. The choice of the right computational tool, done at a early stage of exploratory data analysis, will dictate the generated hypotheses about the underlying biology. Here we demonstrated that Poincaré maps reveal complex cell developmental processes that would remain undiscovered by prior methods. Poincaré maps is able to do so by leveraging hyperbolic geometry and placing minimal assumptions about the data. While any hypothesis generated via computational analysis should be validated in the lab before being converted into strong statements, a properly chosen computational tool will facilitate the selection of appropriate experiments.

For this purpose, Poincaré maps aids the discovery of complex hierarchies from single-cell data by embedding large-scale cell measurements in a two-dimensional Poincaré disk. The resulting embeddings are easy to interpret during exploratory analysis and provide a faithful representation of similarities in hierarchies due to the underlying geometry. This property makes Poincaré maps stand out among other embeddings as it allows to simultaneously handle visualization, clustering, lineage detection, and pseudo-time inference. Poincaré maps do not need to be constrained to two dimensions, and would have the same implementation for 3 or dimensions. However, for the datasets used in this study two dimensions were sufficient; using more dimensions would reduce readability and harden interpretation.

Since Poincaré maps involves several hyper-parameters and non-convex optimization, we thoroughly studied sensitivity the method performance to these parameters. Similar to most manifold learning methods, the number of nearest neighbors *k* will have significant effect on the performance of the method. The tuning of additional hyper-parameters such as *σ* and *γ* will have some small effect on the method’s performance in terms of local and global structure, and are typically easy to select using visual inspection or the scale-independent quality measure. Finally, we observed that the choice of random seed had no significant effect on the visualization properties.

With Poincaré maps, we hope to bring interest about hyperbolic embeddings to the biology community. Due to their advantageous properties for modeling hierarchical data, they could provide substantial benefits for a wide variety of problems such as studying transcriptional heterogeneity and lineage development in cancer from single-cell RNA and DNA sequencing data, reconstructing the developmental hierarchy of blood development, and reconstructing embryogenesis branching trajectories. We also would like to stress that Poincaré maps could be a good candidate embedding to visualize cell atlases of whole organisms as they are able to preserve global similarities between measurements. Finally, we note that Poincaré maps are not limited to the analysis of scRNAseq, but could be applied to any type of data with a hidden hierarchical structure.

## Methods

In the following we discuss the main stages of our method, i.e., estimating proximities that are informative about hierarchical structure and embedding these proximities into the Poincare disk.

Let 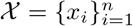 be a high-dimensional dataset of *n* samples ***x***_*i*_ ∈ ℝ^*p*^ (e.g., individual cells) with *p* features (e.g., gene expression measurements).

### Local Connectivity

We first estimate local connectivity structures as typically done in manifold learning^26;28;29^. When the dimensionality of data exceeds 200 dimensions, we suggest to operate on the top 50–100 PCA components. In particular, let 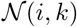 denote the *k* nearest neighbors of ***x***_*i*_ in 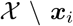 according to the Euclidean distance. We then create a symmetric *k*-nearest-neighbor graph *G* = (*V, E, w*), where the set of vertices 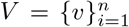 represents the samples in 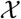 and the set of edges 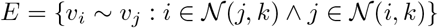 represent the nearest neighbor relations. Furthermore, each nearest neighbor relation is weighted using the Gaussian kernel over distances

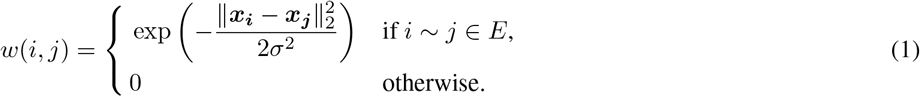

where *σ* is a hyperparameter that controls the kernel width. By enforcing connectivity of *G*, we preserve finite distances between all measurements.

### Global Proximites

To estimate the underlying manifold structure from distances on the *k*NN graph *G*, we can employ all-pairs shortest paths or related methods such as the *Relative Forest Accessibility* (RFA) index, which is defined as follows: Let *L* = *D* − *A* denote the graph Laplacian of the graph *G*, where *A*_*ij*_ = *w*(*i*, *j*) is the corresponding adjacency matrix and *D*_*ii*_ = ∑_*j*_ *w*(*i*, *j*) is the degree matrix. The RFA matrix *P* is then given as^27^

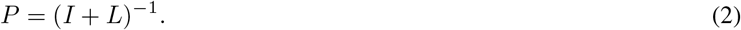

*P* is a doubly stochastic matrix where each entry *p*_*ij*_ corresponds to the probability that a spanning forest of *G* includes a tree rooted at *i* which also includes *j* (i.e., where *j* is accessible from *i*)^27;35^ Compared to shortest-paths, the RFA index has the advantage to increase the similarity between nodes that belong to many shortest paths. This can provide an important signal to discover hierarchical structures as nodes that participate in many shortest paths are likely close to the root of the hierarchy. In all experiments, we use the RFA index to estimate global proximities.

### Hyperbolic Embedding

Given *P*, we aim at finding an embedding ***y***_***i***_ of each ***x***_*i*_ that highlights the hierarchical relationships between the samples. For this purpose, we embed *P* into two-dimensional hyperbolic space.

The Poincaré disk is the Riemannian manifold 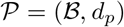, where 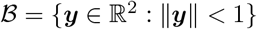 is the open 2-dimensional unit ball. The distance function on 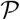 is then defined as

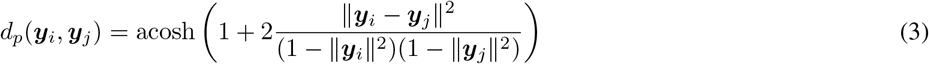

It can be seen from Equation (3), that the Euclidean distance within 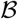 is amplified smoothly with respect to the norm of ***y***_*i*_ and ***y***_*j*_. This property of the distance is key for learning continuous embeddings of hierarchies. For instance, by placing the root node of a tree at the origin of 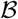, it would have relatively small distance to all other nodes, as its norm is zero. On the other hand, leaf nodes can be placed close to the boundary of the ball, as the distance between points grows quickly with a norm close to one.

To compute the embedding we use an approach similar to t-SNE^17^ and approximate the RFA probabilities in *P* via distances in the embedding space. In particular, we define the similarity *q*_*ij*_ between the embeddings *ν*_*i*_ and *ν*_*j*_ as:

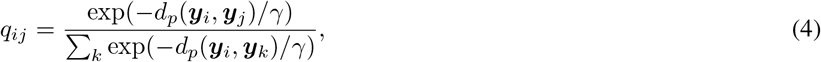

where 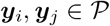. A natural measure for the quality of the embedding is then the symmetric Kullback-Leibler divergence between both probability distributions:

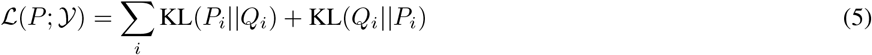

### Details on the optimization

To compute the embeddings, we minimize Equation (5) via *Riemannian Stochastic Gradient Descent* (RSGD).^36^ In particular, we update the embedding of ***y***_*i*_ in epoch *t* using

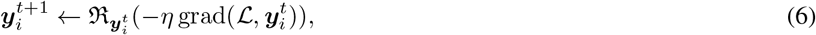

where grad 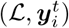 denotes the Riemannian gradient of Equation (5) with respect to 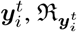 denotes a retraction (or the exponential map) from the tangent space of 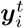 onto 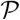, and *η* > 0 denotes the learning rate. The optimization can be performed directly in the Poincaré ball or, alternatively, in the Lorentz model of hyperbolic space which provides improved the numerical properties and efficient computation of the exponential map.^37^

### Translation in 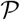

Equation (5) favors embeddings where nodes with short distances to all other nodes are placed close to the origin of the disk. While such nodes correspond often to nodes that are close to the root of the underlying tree, it is not guaranteed that the root is the closest embedding to the origin. However, when the root node is known, we can perform an isometric transformation of the entire embedding that places this node at the origin and preserves all distances between the points. In particular, to translate the disk such that the origin of the Poincaré disk is translated to ***ν***, ***x*** is translated to

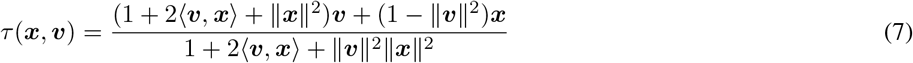

Since the spatial resolution is amplified close to the origin of the disk, provides also a method to “zoom” into different parts of the embedding by moving the area of interest to the origin.

### Clustering

Hyperbolic space is a metric space and thus allows us to compute distances between any pair of points. This makes Poincaré maps straightforwardly applicable to clustering techniques that rely only on pairwise (dis)similarity measurements such as spectral clustering, agglomerative clustering and kmedoids.

### Lineages

As a naive approach for lineage detection we suggest to use agglomerative clustering by the angle between a pair of points in the Poincaré disk after the rotation with respect to the root node.

### Poincaré pseudotime

“Pseudotime” is typically reffered as “a measure of how much progress an individual cell has made through a process such as cell differentiation”^25^. As Poincaré pseudotime we propose to use the distance from the root node in the Poincaré ball.

### Choice of Hyperparameters

In the following, we discuss the function of different hyperparameters in Poincaré maps and propose typical value ranges. The number of nearest neighbors *k* reflects the average connectivity of the clusters and is typically set to *k* ∈ [15, 30]. The Gaussian kernel width *σ* is responsible for the weights for the *k*-NN graph in the original space and is typically set to *σ* ∈ [1.0, 2.0]. The softmax temperature *γ* controls the scattering of embeddings and is typically set to *γ* ∈ [1.0, 2.0].

### Scale-independent quality measure

To quantitatively compare performance of different embedding approaches, we use the scale-independent quality criteria proposed by Lee et al.^34^ The main idea of this approach is that a good dimensionality reduction approach, will have a good preservation of local and global distances on the manifold, e.g. close neighbours should be placed close to each other, while maintaining large distances between distant points. Lee et al.^34^ proposed to use two scalar quality criteria *Q*_*local*_ and *Q*_*global*_ focusing separately on low and hight dimensional qualities of the embedding. The quantities of *Q*_*local*_ and *Q*_*global*_ range from 0 (bad) to 1 (good) and reflect how well are local and global properties of the dataset are preserved in the embedding (see details in **Supplementary Note 2**). To estimate distances in the high-dimensional space *δ*_*ij*_, we use geodesic distances estimated as length of a shortest-path in a *k*-nearest neighbours graph. We fixed *k* = 20 for all the datasets as there is no objective way to decide on a correct *k* and visually results looked good for all the embeddings for this choice of *k*. For the distances low-dimensional space we use euclidean space for all the embeddings except Poincaré maps, for which we use hyperbolic distances. As all the embeddings involve an element of stoachasitcity in their output, we run every embedding three times with a different seed. We run all the embeddings with a different set of parameters.

### Computational complexity and time

Memory complexity of Poincaré maps is *O*(*n*^2^), where n is the number of samples. Time complexity consits of three parts: estimation of *k*NNG − *O*(*n*^2^) (this part could be replaced with FAISS^38^ for scalability), estimation of RFA − *O*(*n*^2^) and minimization of KL-divergence – *O*(*neb*), where e - maximum number of epochs, *b* - batch size. As we need to minimize KL till convergence, we can in advance estimate the number of epochs needed. For all the datasets used here the number of epochs was less than 2000 and we also used early stopping upon convergence. Typical running time on 1 GPU for all the small-medium datasets is less than a minute, and for large datasets around 15 min (Planaria) or 2-3 hours (C. elegans).

## Code availability

The code is available at https://github.com/facebookresearch/PoincareMaps.

## Data availability

Several public datasets were used in this study: three synthetic datasets generated with Scanpy, Olsson et al. (synapse ID syn4975060), Paul et al. (accession code GSE72857), Moignard et al. (accession code GSE61470), Plass et al. (accession code GSE103633, preprocessed data available at https://shiny.mdc-berlin.de/psca/), Packer et al. (preprocessed data available at https://github.com/qinzhu/VisCello).

## Acknowledgments

We would like to thank Ioana Sandu and Will Macnair for valuable discussions.

## Author information

### Contributions

L.B., M.N. and A.K. conceived the idea. A.K., M.N. and D. L.-P. designed and implemented the computational tools. A.K. performed the analysis and biological interpretation of results. L.B. contributed to the design of the study. M.N. supervised the study. A.K., M.N. and D.L.-P. wrote the manuscript.

### Corresponding author

Correspondence to Anna Klimovskaia or Maximilian Nickel.

## Supplementary Notes

### Supplementary Note 1: Poincaré maps for learning hierarchical representations

Hyperbolic space is a Riemannian manifold whose structure is well-suited to represent hierarchical and tree-like relationships. For our work, this combines two important advantages: First, the metric structure of hyperbolic space allows us to capture continuous hierarchical relationships and interpolate between points. Second – and in contrast to other metric spaces – hierarchies can already be represented in two-dimensional hyperbolic space with small distortion [1, 2, 3, 4].

#### Poincaré disk model

There exist multiple, equivalent models of hyperbolic space, such as the Beltrami-Klein, the Lorentz, and the Poincaré half-plane model. In this work, we base our approach on the Poincaré disk model, as it is best suited for visual analysis. The Poincaré disk is defined as follows: let 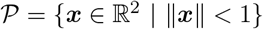 be the *open* unit disk, where ||·|| denotes the Euclidean norm. The Poincaré disk corresponds then to the Riemannian manifold 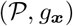, i.e., the open unit disk equipped with the Riemannian metric tensor

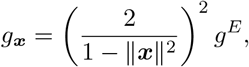

where 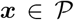 and *g*^*E*^ denotes the Euclidean metric tensor. Furthermore, the distance between points 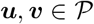 is given as

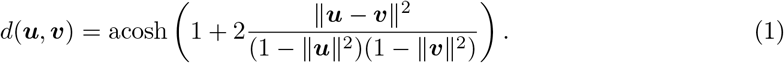

**Supplementary Figure 1.**
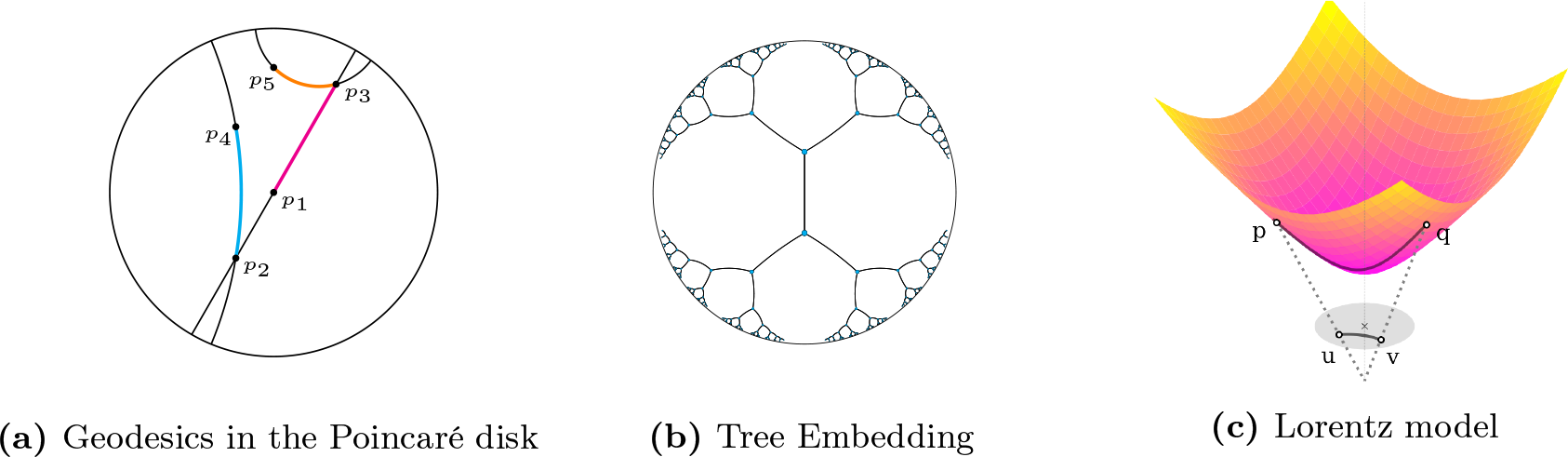
a) Geodesics in the Poincaré disk model of hyperbolic space. Due to the negative curvature of the space, geodesics between points are arcs perpendicular to the boundary of the disk. For curved arcs, midpoints are closer to the origin of the disk (p1) than the associated points, e.g. (p3, p5). c) Points (p,q) lie on the surface of the upper sheet of a two-sheeted hyperboloid. Mapping of points on the hyperboloid (p, q) onto the Poincaré disk.

The boundary of the disk is denoted by 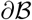 and is not part of the manifold, but represents infinitely distant points. Geodesics in 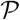 are then arcs orthogonal to 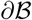 (as well as all diameters). See Figure 1a for an illustration.

It can be seen from Equation (1) that the Euclidean distance of two points in the Poincaré disk is amplified with respect to their distance to the origin of the disk. This locality property of the Poincaré distance is key for continuous embeddings of hierarchies. For instance, by placing the root node of a tree at the origin of 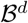 it would have a relatively small distance to all other nodes as its Euclidean norm is zero. On the other hand, leaf nodes can be placed close to the boundary of the Poincaré disk, as the distance grows fast between points with a norm close to one. Furthermore, Equation (1) is symmetric and the hierarchical organization of the space is solely determined by the distance of points to the origin. Due to this property, Equation (1) is applicable in an unsupervised setting, where the hierarchical order of objects is not specified in advance. Importantly, this allows to learn embeddings that simultaneously capture the hierarchy of objects (through their norm) as well as their similarity (through their distance).

The Riemannian manifold structure of hyperbolic spaces enables the use Riemannian Stochastic Gradient Descent (RSGD) [5] to compute the embeddings. In RSGD, parameter updates are performed via

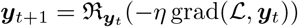

where 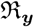 denotes a retraction from the tangent space at ***y*** onto the manifold, grad 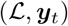 denotes the Riemannian gradient of the scalar function 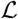, and *η* > 0 denotes the learning rate. The embeddings can be learned directly in the Poincaré disk 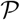 or, alternatively, in the Lorentz model (which has advantageous properties for stochastic optimization). We refer to [2] and [4] for the detailed optimization procedure on both hyperbolic manifolds. When optimization is performed in the Lorentz model, we can map the learned embeddings into the Poincaré disk via the diffeomorphism 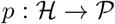, where

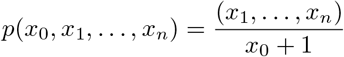

which preserves all geometric properties including isometry (see also **Supplementary Figure 1**).

### Supplementary Note 2: Benchmarks on datasets with known hierarchy

#### Visualization

We compare Poincaré maps to several methods frequently used for visualization: tSNE [6], UMAP [7], diffusion maps [8], graph abstractions (PAGA) [9], ForceAtlas2 [10] and Monocle 2 [11]. For all competing methods, we used a set of parameters in the range provided by the authors. For the visualization comparison, for each method we chose the best set of parameters in terms of quality metric described below.

While methods such as diffusion maps, PAGA and Monocle 2 can be used by a knowledgeable user to infer the correct structure form data with several post-processing iterations, here we would like to demonstrate how Poincaré maps extracts meaningful insights from data without further post-processing. The ability to recover hidden hierarchies automatically and in one shot makes Poincaré maps an attractive tool for the analysis of branching processes and complex hierarchical structures.

#### Scale-independent quality criteria

To quantitatively compare the performance of different embedding approaches, we use a scale-independent quality criteria proposed by Lee et al. [12] The main idea is that a good dimensionality reduction approach will have a good preservation of local and global distances on the manifold, e.g. close neighbors should be placed close to each other, while maintaining large distances between distant points. Below we provide a short summary of how to compute this metric. All the details can be found in the original paper of Lee et al.[12]

Let 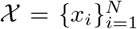 be a high-dimensional dataset of *N* samples ***x***_*i*_ ∈ *ℝ*^*p*^ (e.g., individual cells) with *p* features (e.g., gene expression measurements) and 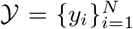 be a low-dimensional representation of this dataset in *m* = 2 dimensions. Let *δ*_*ij*_ denote the distance from *x*_*i*_ to *x*_*j*_ in the high-dimensional space and *d*_*ij*_ denote the distance from *y*_*i*_ to *y*_*j*_ in the low-dimensional space. Assume *δ*_*ij*_ = *δ*_*ji*_ and *δ*_*ij*_ = *d*_*ji*_. The distances could be used to compute high (*R* = {*ρ*_*ij*_}_1≤ *i*,*j*≤*N*_) and low (*V* = {*v*_*ij*_}_1≤ *i*,*j*≤*N*_) dimensional ranks between the points:

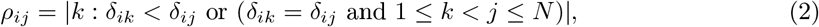

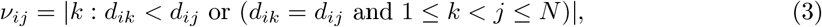

where |*A*| denotes the cardinality of a set. According to this definition, reflexive ranks are set to zero and non-reflexive ranks belong to {1,…, *N* − 1}.

A co-ranking matrix **Q** = {*q_kl_*}_1≤*k*,*l*≤*N* − 1_ is defined as:

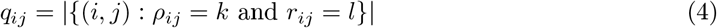

The co-ranking matrix contains all the necessary information about how ranks are preserved in a given low-dimensional representation. As was demonstrated by Lee et al. [12], the co-ranking matrix **Q** is straightforward to compute, and it could be used to compute *Q*_*NX*_ – scale-independent quality criteria for dimensionality reduction for a given value of *K* = 1,…, *N* − 1:

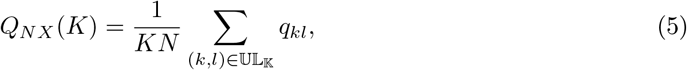

where 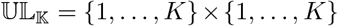 is the upper left corner of co-ranking matrix. *Q*_*NX*_(*K*) ∈ [0, 1] assesses the overall quality of the embedding. Essentially, it measures the preservation of K-ary neighborhoods. A perfect embedding has *Q*_*NX*_(*K*) = 1 for every *K* = 1,…, *N* − 1.

The left part of of the *Q*_*NX*_(*K*) curve reflects how local properties are preserved, and the right part corresponds to the preservation of global properties. To improve its readability, Lee et al.[12] propose to use two scalar quality criteria *Q*_*local*_ and *Q*_*global*_ focusing separately on low and hight dimensional qualities of the embedding:

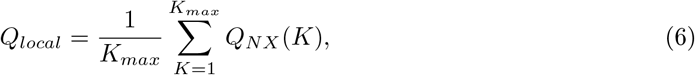

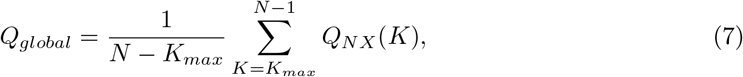

where *K*_*max*_ defines the split of the *Q*_*NX*_ curve and is automatically computed as:

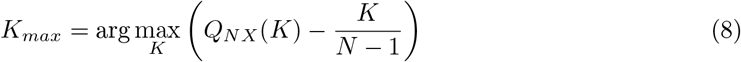

The quantities of *Q*_*local*_ and *Q*_*global*_ range from 0 (bad) to 1 (good).

In this work, to estimate distances *δ*_*ij*_ in the high-dimensional space, we use geodesic distances estimated as the length of a shortest-path in a *k*-nearest neighbors graph. We fixed *k* = 20 for all the datasets, as there is no objective way to decide on a correct *k,* and visually results looked good for all the embeddings for this choice of *k*. For the distances *δ*_*ij*_ in the low-dimensional space we use euclidean space for all the embeddings except Poincaré maps, for which we use hyperbolic distances. As all the embeddings involve an element of stochasticity in their output, we run every embedding three times with a different seed. We run all the embeddings with a different set of parameters in the range proposed by the authors of each method. **Supplementary Figure 2** demonstrates the comparison of *Q*_*local*_ and *Q*_*global*_ for all the datasets described below.

### Robustness of Poincaré maps to random seed and choice of hyper-parameters choice

We used the quality criteria described above and visual inspection to address the robustness of Poincaré maps to hyper-parameters choice and random seed. **Supplementary Figure 3 (a)** demonstrates that good values of *σ* vary for different datasets. However, the parameter *γ* has a less strong effect and rather controls how much the embedding will be scattered on the disk. We advise to set *γ* to 1.0 or 2.0 depending on the dataset size: larger datasets typically have better visualization with *γ* = 2.0 (**Supplementary Figure 3 (b-c)**). **Supplementary Figure 3 (a, d)** demonstrates that Poincaré maps are very robust to random seed and that it doesn’t significantly affect neither quality, nor visual interpretation.

**Supplementary Figure 2.**
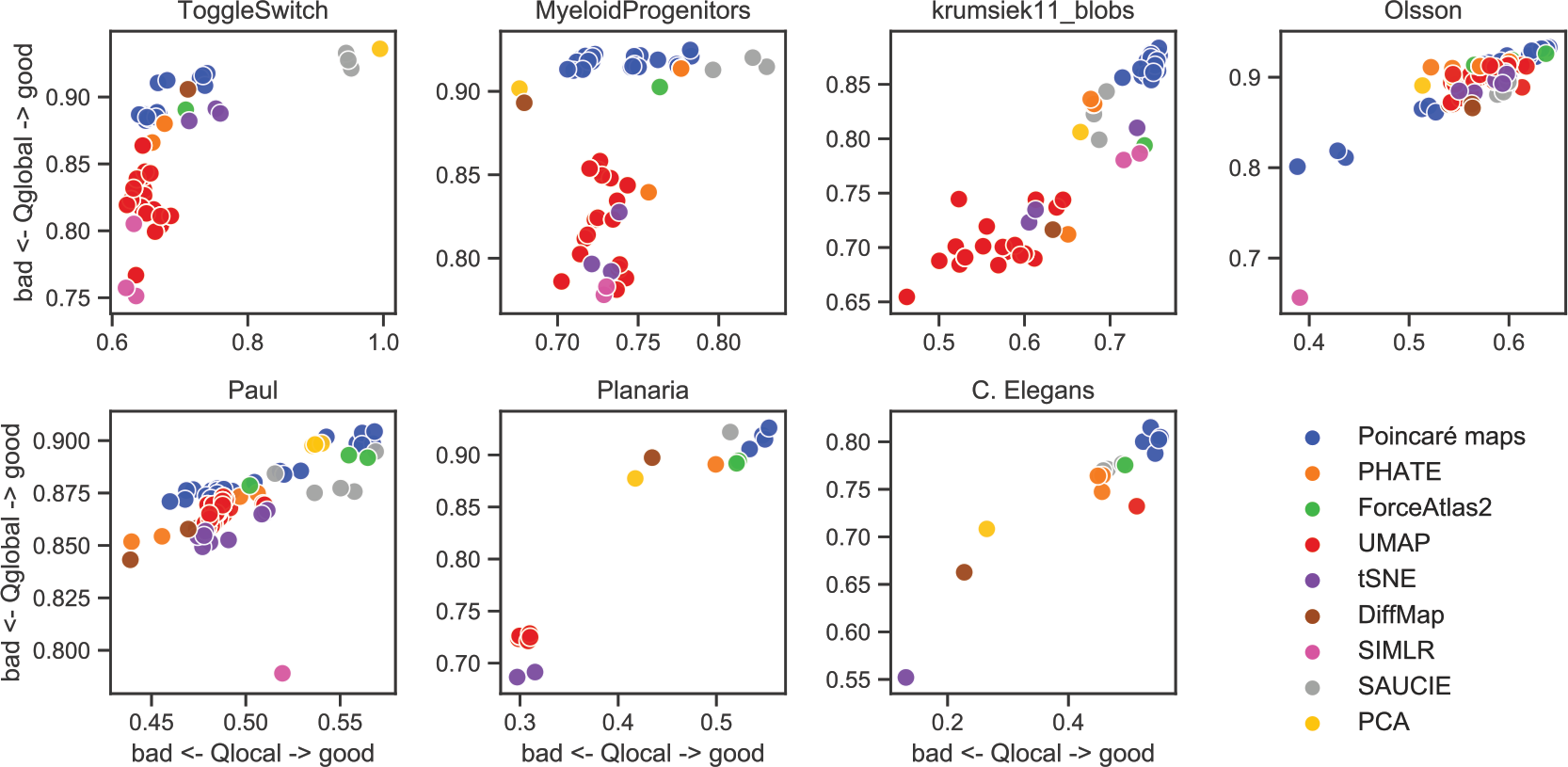
Comparison of local and global quality metrics for various datasets. All embeddings were computed with 3 random seeds and several different hyper-parameters. We fixed *k* = 20 for all the datasets for a fair comparison.

#### Synthetic datasets

To demonstrate the performance of Poincaré maps we used several synthetic datasets available as Jupyter notebooks with Scanpy [13]: a simple toggle switch, myeloid progenitors and myeloid progenitors with Gaussian blobs. These datasets were previously used to demonstrate the performance of diffusion maps and graph abstractions, and constitute great examples of manifolds with a hierarchical structure of increasing levels of complexity. All models consist of Boolean equations, which were translated into ordinary differential equations and simulated with Scanpy as stochastic differential equations with Gaussian noise [14].

A simple toggle switch model [15, 16] is a process with two branches, which are defined by the expression of two markers. **Supplementary Figure 4** demonstrates that all competing methods produce rather correct results for this simple problem. However, Poincaré maps gives a more clear separation of the intermediate states of terminal fates (inter1 vs inter2). In this example, only tSNE, diffusion maps, and Poincaré maps produce embeddings with meaningful pairwise distances.

A synthetic dataset for myeloid differentiation [17] represents cell differentiation progresses of a common myeloid progenitor state towards one of four different branches: erythrocyte, neutrophil, monocyte and megakaryocyte. **Supplementary Figure 5** shows the provided embeddings for all methods. Poincaré maps produce an embedding which is visually similar to the other methods, but has neither discontinuities, nor overlaps in the trajectories, since it preserves all the pairwise distances. Given the known root, the rotation of the Poincaré map (by means of translation) allows to easily read out the hierarchy. Diffusion maps produce embeddings consistent with one main branch, but more Euclidean dimensions would be necessary to separate the rest. Monocle 2 produce a tree layout consistent with the hierarchical structure of the data, but is not able to reconstruct the temporal connection (trajectory) of the cell differentiation process.

The third dataset shows the stability of Poincaré maps with respect to the existence of clusters not related to the main cell development process. To this end we use the synthetic dataset of myeloid differentiation with two Gaussian blobs, added as proposed by Wolf et al. [9] (**Supplementary Figure 6**). None of the benchmark methods except ForceAtlas2 is able to capture the hierarchy.

#### Mouse myelopoesis dataset (single-cell RNA seq)

To demonstrate the performance of Poincaré maps on single-cell RNA seq data, we used the mouse myelopoesis dataset (wild type only) from Olsson et al. [18]. The data was downloaded and preprocessed according to the pipeline from Qiu et al. [11]. The processed dataset contained 532 features for 382 cells. Nine cell types were annotated corresponding to the original study: HSCP-1 (haematopoietic stem cell progenitor), HSCP-2, megakaryocytic, erythrocytic, Multi-Lin* (multi-lineage primed), MDP (monocyte-dendritic cell precursor), monocytic, granulocytic and myelocyte (myelocytes and metamyelocytes). In order to obtain the best results for Monocle 2, we used the analysis pipeline provided by the authors (https://github.com/cole-trapnell-lab/monocle2-rge-paper). As the reference hierarchy, we used the canonical hematopoetic cell lineage tree [19] (**Supplementary Figure 7 (a)**).

**Supplementary Figure 3.**
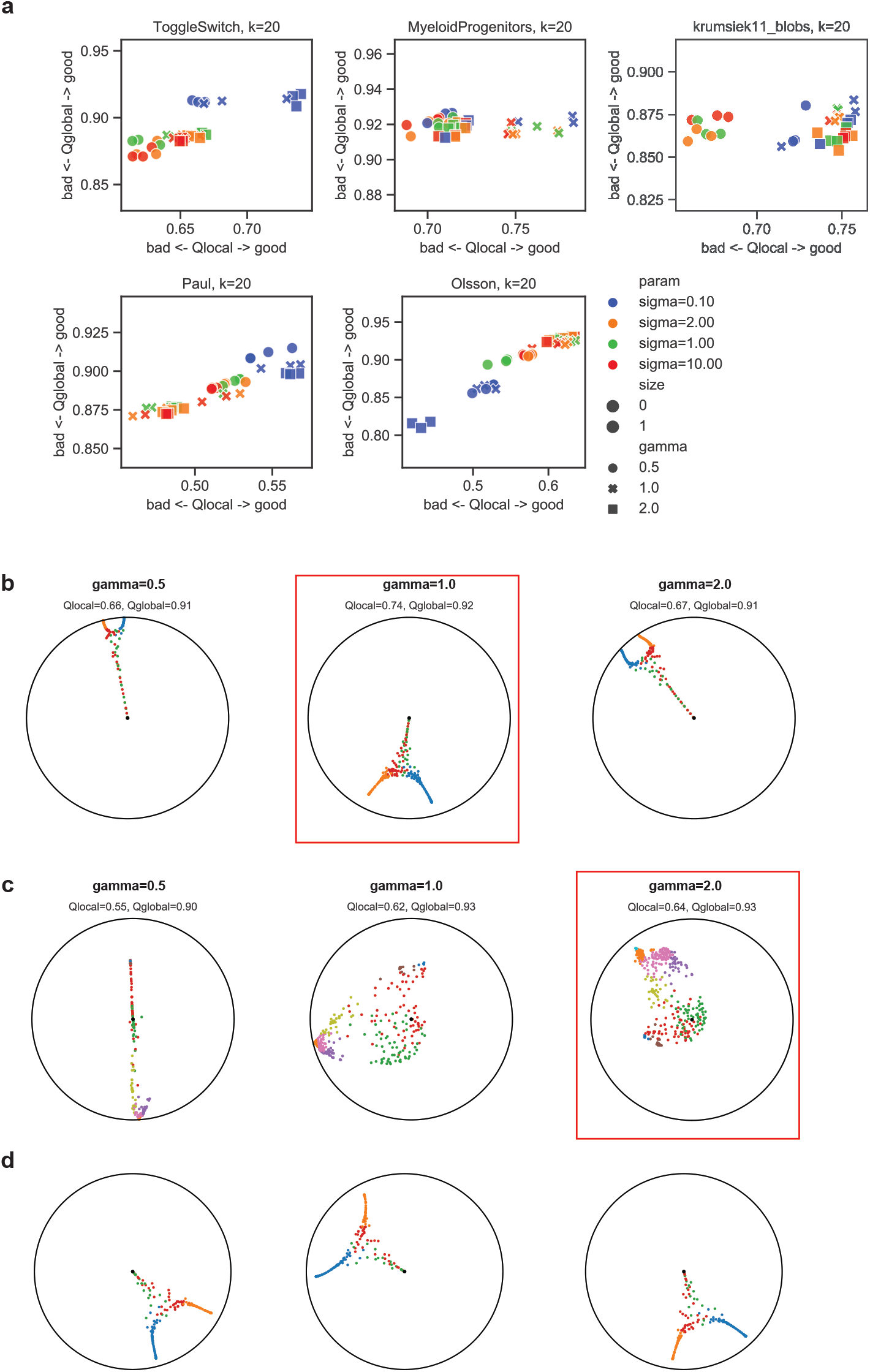
Robustness of Poincaré maps to random seed and hyper-parameters choice for a fixed. *k* = 20 **(a)** Scale-independent quality criteria for various hyper-parameters and three random seeds. **(b – c)** Change of visual qualities of the embedding for a fixed *σ* and varying *γ* for ToggleSwitch (b) and Olsson (c) datasets. Red frame represents best quality score. **(d)** Comparison of robustness to random seed: *σ* and *γ* are fixed.

Poincaré maps, after rotation (**Supplementary Figure 7 (b)**), reveal the known hierarchy and suggest that part of HSPC-2 cluster actually corresponds to the megakaryocyte/erythrocyte progenitor (MEP), and that the cluster named Multi-Lin corresponds to the granylocyte/monocyte progenitor (GMP). Also according to Poincaré maps, the cluster annotated as myelocyte does not belong to the hierarchy, or constitutes a mature state of granulocytes. However, the validation of these hypotheses requires a detailed differential expression analysis.

**Supplementary Figure 7 (c)** shows how widely used methods such as tSNE distort the pairwise distances, therefore making more difficult to draw conclusions about hierarchies. Similarly, two dimensions of diffusion maps are not enough to represent the branching. UMAP and ForceAtlas2 results overall agree with the Poincaré maps, but don’t allow to reason about the subtle hierarchical relations between HSCP-1/2 clusters and MDP. Monocle 2 captures the global branching, but fails to depict more fine-grained relations: between erythrocytytes and megakaryocytes or granulocytes and myelocytes.

#### Mouse myeloid progenitors dataset (MARS-Seq)

As an example of a dataset with multiple intermediate populations, we use a dataset provided by Paul et al. [20]. Myeloid progenitor cells were separated by sorting the c-Kit+ Sca1 lineage from mouse bone marrow and sequenced with MARS-seq. We followed the data preprocessing procedure recipe_zheng17 (Scanpy-recipe [21]), which selects the 1000 most highly-variable genes for 2730 cells. In the original study, the authors identify 19 clusters. We use these labels and canonical hematopoetic cell lineage tree (**Supplementary Figure 8 (a)**) to compare the performance of all methods. We run all methods except Monocle 2 on the 20 top principal components of the preprocessed data. For Monocle 2, we used the Jupyter notebook provided by the authors (the lymphoid cluster was separated as described in the original study).

**Supplementary Figure 8 (b)** shows the embeddings provided by Poincaré maps. For this dataset, the root is supposed to be at CMP cluster, which is not observed. We chose the root as the medoid (with respect to Poincaré distances) of the MEP and GMP clusters combined. **Supplementary Figure 8 (c)** shows the hierarchy that could be read out from the Poincaré map. We would like to point out that Poincaré maps clearly separate lymphoid cells and dendritic cells as outliers, which agrees with the canonical tree as they are part of lymphoid lineage. None of the other methods **(Supplementary Figure 8 (d)**) were able to capture this fact. Poincaré maps also suggest that some of the clusters (13-15) could be relabeled to better reflect the canonical hierarchy. After the removal of the lymphoid cluster, Monocle 2 captures the main lineage branching between the MEP and GMP lineages, but it does not separate dendritic cells, and destroys the eosonphils cluster. Wolf et al. demonstrated that Monocle 2 results without the removal of the lymphoid cluster only worsen.

Finally, Poincaré maps places the 16Neu cluster downstream of 15Mo in the hierarchy. However, the canonical hierarchy shows neutrophils and monocytes at the same level. As noted by Wolf et al., we suppose that this inconsistency is due to a faulty labeling of the clusters.

#### Planaria dataset (Drop-seq)

To demonstrate scalability of Poincaré maps to large datasets, we analyzed the entire Planaria dataset of Plass et al. [22]. The dataset comprises 11 individual experiments capturing a total of 21,612 cells with droplet-based single-cell transcriptomics (Drop-seq). To obtain the Poincaré maps we used the pre-processed data provided by the authors: https://nbviewer.jupyter.org/github/rajewsky-lab/planarian_lineages/blob/master/paga/planaria.ipynb. The preprocessed dataset comes in the form of 50 principal components, which were used by the authors to apply tSNE, PAGA and ForceAtlas2. (**Supplementary Figure 9**) illustrates that Poincaré maps agree with tSNE and ForceAtlas2 embeddings, significantly outperform PCA and UMAP, and agrees with the PAGA hierarchy annotation (Figure 4 in Plass et al.). Unfornutately we were not able to compute SIMLR for this dataset, because of the computation time.

#### C. elegans dataset (10X Genomics)

The C. elegans dataset from Packer et al. [23] is the largest dataset used in our experiments. Original dataset contained 84,625 single cells measured with 10x Genomics platform. We used the preprocessed version (batch corrected, 100 PCA components) of the data provided by the authors at https://github.com/qinzhu/VisCello. In the original study the authors used UMAP to visualize the data. We loaded the UMAP coordinates provided by the authors to perform fair comparison with Poincaré maps. 37 manually annotated labels of cell types were provided together with the dataset. We randomly down-sampled the dataset to 40,000 cells. The whole dataset represents > 60x oversampling of the 1,341 branches in the C. elegans embryonic lineage, therefore down-sampling should not destroy statistical properties of the dataset. We checked that sub-sampled data contained all 37 original cell types. **Supplementary Figures 2, 10, 11** demonstrates, that Poincaré maps significantly outperform all other embedding methods. Unfortunately we were not able to compute SIMLR for this dataset, because of the computation time.

#### Clustering

Poincaré maps provide embeddings useful beyond visualization. Since Poincaré maps preserve pairwise similarities, their embeddings are suitable for downstream tasks, such as clustering. We compared several clustering approached using Poincaré maps and benchmark embeddings. We also provide Louvain clustering and clustering in the original gene expression space. Since the datasets comprise several continuous trajectories and there is no true separation for progenitor populations of different branches, we used the Adjusted Rand Index (ARI) and Fowlkes-Mallows scores (FMS) to measure cluster quality.

##### Adjusted Rand Index

The Adjusted Rand Index (ARI) is a function that measures the similarity between two cluster assignments. ARI is bonded bewteen [−1, 1], where negative values correspond to independent labelings, similar clusterings have a positive ARI, and 1.0 is the perfect match score. Lets denote *C* a ground truth class assignment and *K* the clustering. Adjusted Rand Index is defined through raw Rand Index (RI):

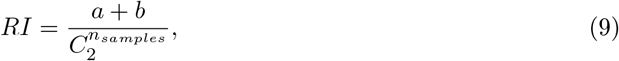

where *a* is the number of pairs of elements that are in the same set in *C* and in the same set in *K*, *b* is the number of pairs of elements that are in different sets in *C* and in different sets in *K*, and 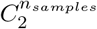 is the total number of possible pairs in the dataset (without ordering). ARI is after adjusting for random labelings:

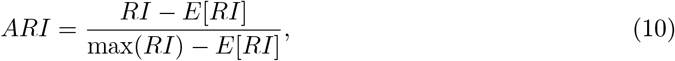

where *a* is the number of pairs of elements belonging to the *same* cluster in predicted and true labels, *b* is the number of pairs of elements belonging to *different* clusters in predicted and true labels, and 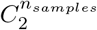 is the number of all possible combinations of pairs of elements in the dataset.

##### Fowlkes-Mallows scores

The Fowlkes-Mallows score FMI is defined as the geometric mean of the pairwise precision and recall:

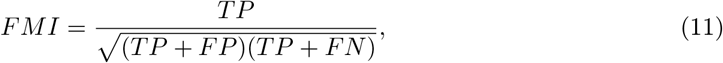

where TP is the number of pairs of points that belong to the same cluster in both the true labels and the predicted labels (true positives), FP is the number of pairs of points that belong to the same clusters in the true labels but not in the predicted labels (false positives), and FN is the number of pairs of points that belong in the same clusters in the predicted labels but not in the true labels (false negatives). The FMI ranges from 0 to 1. A high value indicates a good similarity between two clusterings.

*More details on these metrics can be found at* : https://scikit-learn.org/stable/modules/clustering.html#clustering-evaluation

**Supplementary Table 1** shows the clustering results on synthetic datasets. Poincaré maps achieve very similar score to Louvain clustering, and significantly outperform clustering approaches using other embedding methods, except tSNE embedding, which combined with spectral clustering achieves the best scores. However, as we demonstrated before, tSNE does not preserving the hierarchy, and therefore would be less useful for other downstream tasks.

#### Pseudotime

We demonstrated Poincaré pseudotime performance by comparison with real time and diffusion pseudotime on synthetic datasets. **Supplementary Table 2** demonstrates that Poincaré pseudotime as well as diffusion pseudotime achieve high correlation scores with actual time on all synthetic datasets. This is unsurprising, since these two measures are related in their nature. The performance of both pseudotime approaches is probably bounded by the construction of *k*NNG.

##### Technical details

In the myeloid progenitors dataset, to obtain a pair of points between which to draw the interpolation, we first perform a Louvain clustering of the shrinked dataset. Afterwards, we select two clusters from which we sample a corresponding pair: one cluster corresponding to mature neutrophils, and another cluster corresponding to early neutrophil progenitors.

In the Olsson et al. dataset, we use the previously annotated clusters, and randomly sample several pairs of points belonging to “HSPC-1” cluster, and to either the “Multi-lin”, “Eryth” and “Meg” clusters. This corresponds to a potential position of “HSPC-2” cluster in the hierarchy.

In the Plass et al. Planaria dataset, we removed parenchymal progenitors. We interpolate between the original cluster of parenchymal progenitors and either the “psap+ parenchymal cells”, “pgrn+ parenchymal cells”, or “ldlrr-1+ parenchymal cells”.

As for the network architecture, we used 5 fully connected layers with ReLU non-linearities. For UMAP and ForceAtlas2, we added batch normalization, as it showed better performance.

### Supplementary Note 3: Reconstructing developmental trajectories of asynchronous process: early blood development in mice (qPCR)

We analyze the single cell qPCR dataset of early blood development in mice [24] using Poincaré maps. We followed the data preprocessing procedure described in Haghverdi et al. [8].

First, we visualized the dataset with a Poincaré map using the labels corresponding to different stages of differentiation [24]: primitive streak (PS), neural plate (NP), head fold (HF), four somite GFP negative (4SG−) and four somite GFP positive (4SG+) (**Supplementary Figure 12 (a)**). We see one cluster standing out. Therefore, we perform spectral clustering with Poincaré distances to break down this cluster for further analysis (**Supplementary Figure 12(b),(c)**). Then, cluster 4 mainly consists of Flk1-Runx1-cells (see **Supplementary Figure 12 (d)**). Moignard et al. [24] refer to this cluster as “mesodermal cells at primitive strike” and suggest that these cells give rise to blood and endothelial cells.

The cell that Haghverdi et al. choose as root of the differentiation for the diffusion pseudotime analysis belongs to the “mesodermal” cluster in our analysis. We visualize (**Supplementary Figure 13**) the diffusion pseudotime and Poincaré pseudotime with the roots (a) suggested by Haghverdi et al., and (b) the most dissimilar point in the PS cluster in terms of Poincaré distance. Undesiredly, the distances from (a) grow orthogonal to the actual developmental stages. It agrees with the conclusion in Haghverdi et al. that such a choice of embedding does not allow to see the asynchronous development. Therefore, cluster 4 may not correspond to cells leading to endothelial and blood cells, but rather to early mesodermal cells, which in their turn lead to some other population (Supplementary Figure 4 in Moignard et al.). We will further refer to the cluster 4 as “mesodermal”.

As pointed out by Moignard et al., blood development is a highly asynchronous process, which is hard to capture with PCA or diffusion maps. In **Supplementary Figure 14** we further demonstrate how Poincaré maps could be used to reveal the developmental structure in this process. First, we apply the rotation to the Poincaré map to place the root cell defined above to the center of the disk. Then, we apply our lineage detection procedure and demonstrate that inside of each lineage, the order of the developmental stages is on average preserved. However, if we look at all lineages combined, then the populations from PS, NP, HF stages appear to be a homogeneous mixture. Therefore, the angular information in Poincaré maps adds the additional amount of information crucial to understand asynchronous processes.

Finally, we analyzed the expression profiles of main endothelial and hematopoetic markers for different lineages (**Supplementary Figure 15**). Poincaré maps suggest that cells make an early decision about which branch to become. In particular, we suggest that cells commit to their future branch as early as in the PS stage.

**Supplementary Figure 4.**
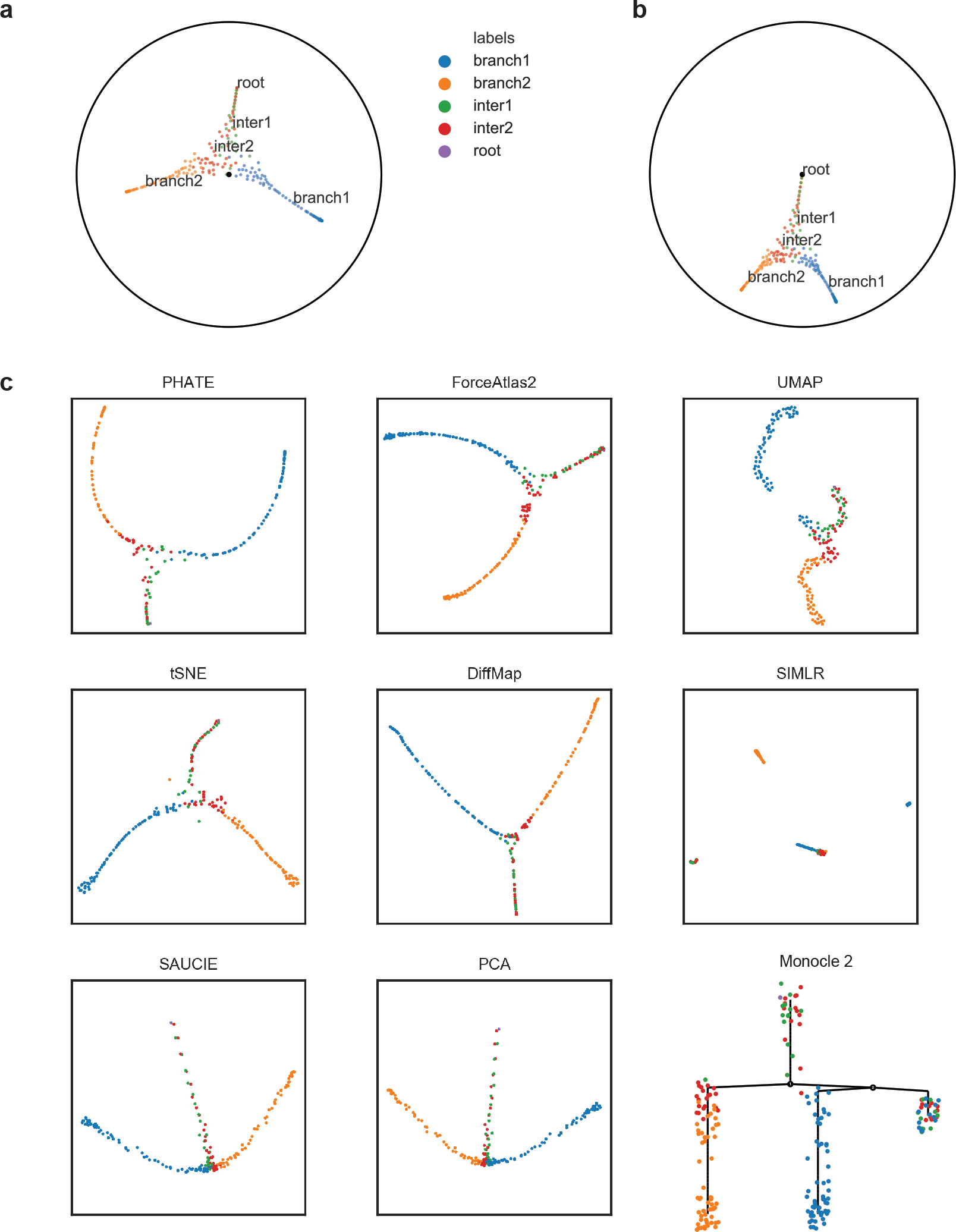
Comparison of various embeddings for the simple toggle switch model. There are two distinct branches. We additionally labeled intermediate states from the simulations. **(a)** Raw Poincaré map. **(b)** Rotation of the Poincaré map with respect to the known root. **(c)** Benchmark methods.

**Supplementary Figure 5.**
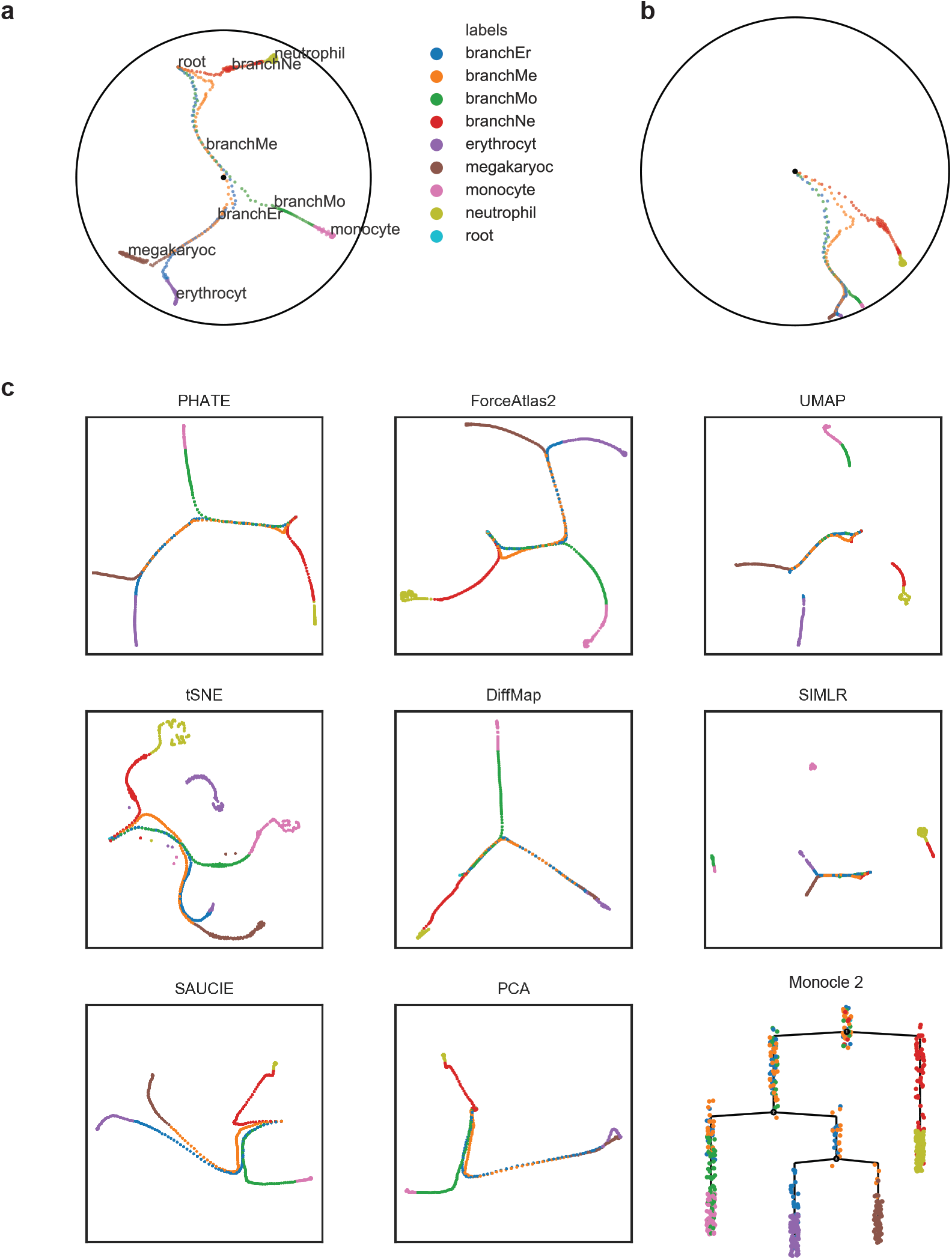
Comparison of various embeddings for a synthetic model of myeloid progenitors differentiation. There are four distinct branches. We additionally labeled intermediate states from the simulations. **(a)** Raw Poincaré map. **(b)** Rotation of the Poincaré map with respect to the known root. **(c)** Benchmark methods.

**Supplementary Figure 6.**
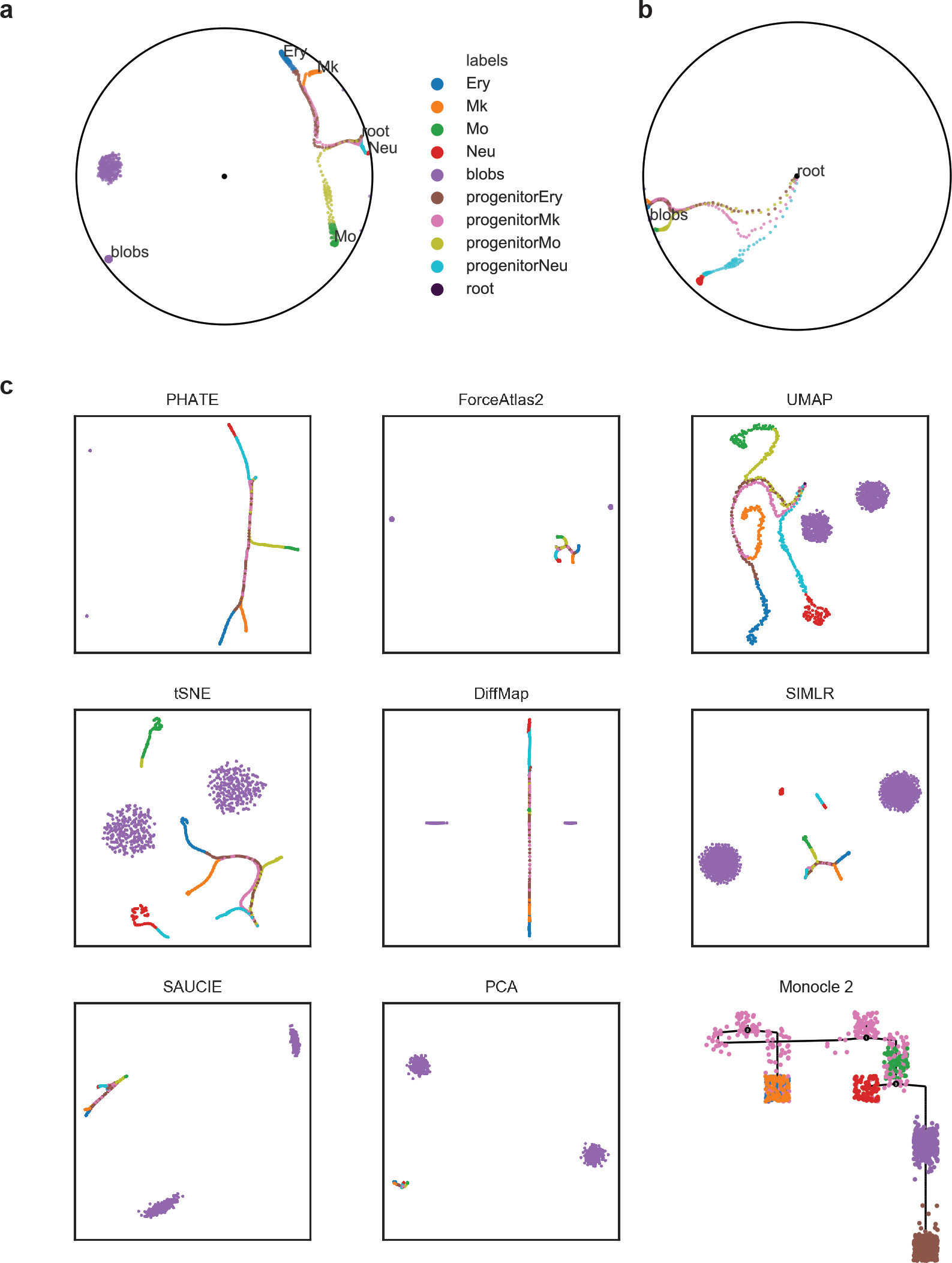
Comparison of various embeddings for a synthetic model of myeloid progenitors differentiation (4 distinct branches) with two additional Gaussian clusters. We additionally labeled intermediate states from the simulations. **(a)** Raw Poincaré map. **(b)** Rotation of the Poincaré map with respect to the known root. **(c)** Benchmark methods.

**Supplementary Figure 7.**
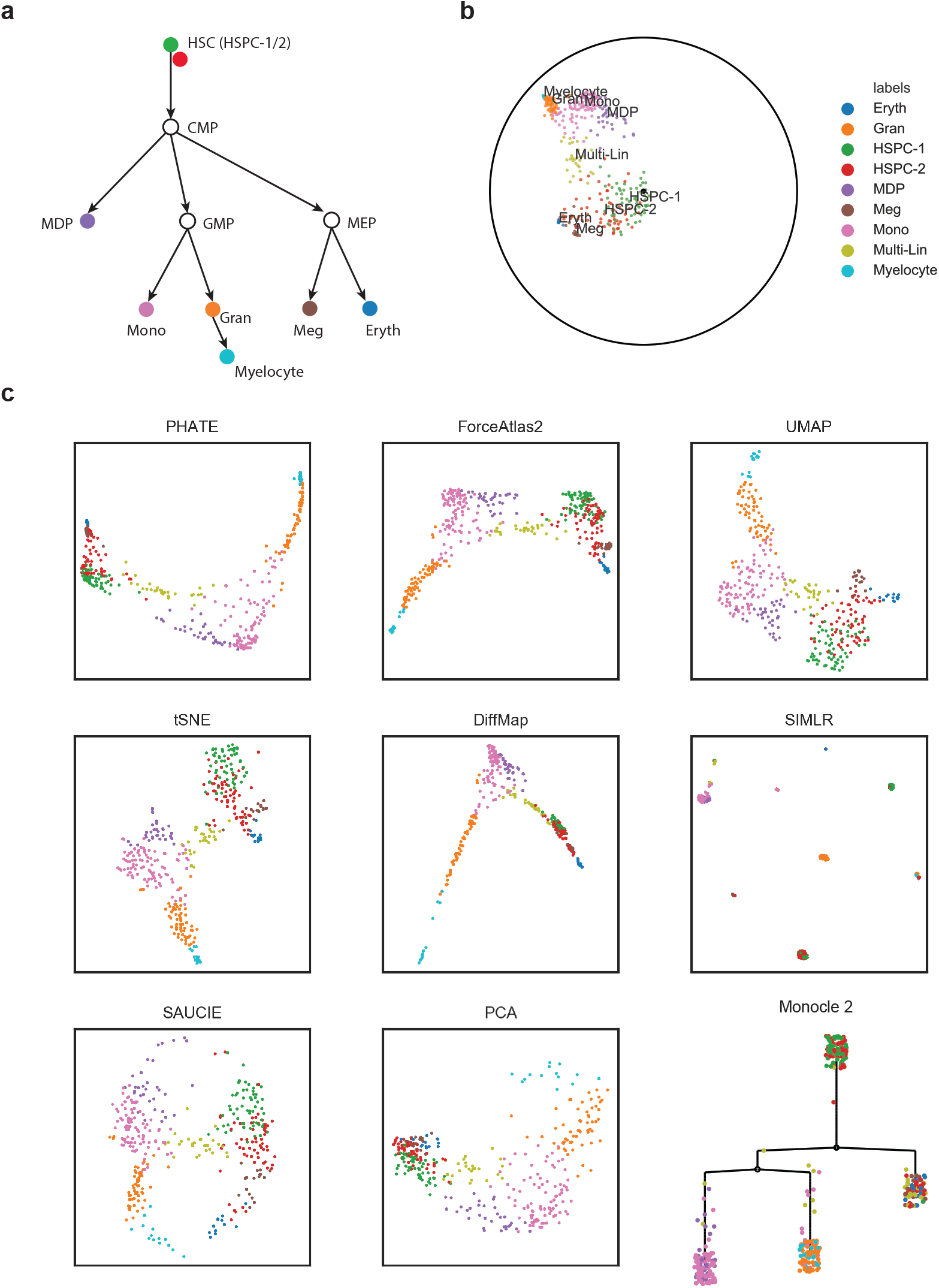
Comparison of various embeddings for the scRNAseq dataset of mouse myelopoesis (Olsson et al.). **(a)** Canonical hematopoetic cell lineage tree. Colored circles correspond to the population colors from the dataset. **(b)** Poincaré map rotated with respect to the root. **(c)** Benchmark methods.

**Supplementary Figure 8.**
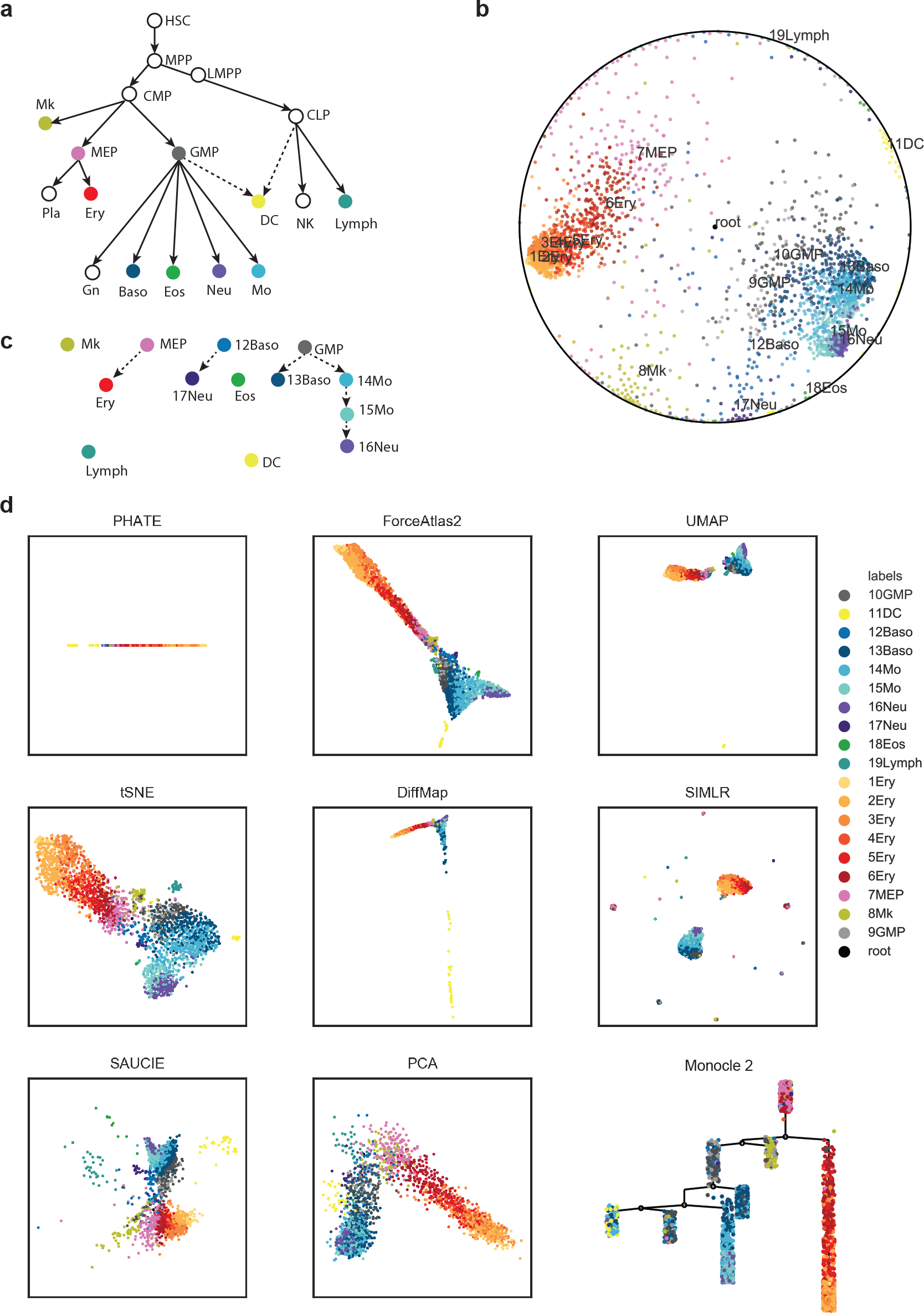
Comparison of various embeddings for the mouse myeloid progenitors MARS-seq dataset (Paul et al.). **(a)** Canonical hematopoetic cell lineage tree. Colored nodes correspond to the population colors from the dataset. White nodes correspond to intermediate annotated states. **(b)** Rotated Poincaré map with respect to the root (medoids of MEP and GMP cluster). **(c)** Hierarchical relationships suggested by the Poincaré map. **(d)** Benchmark methods. To reproduce the Monocle 2 tree, the lymphoid cluster was removed.

**Supplementary Figure 9.**
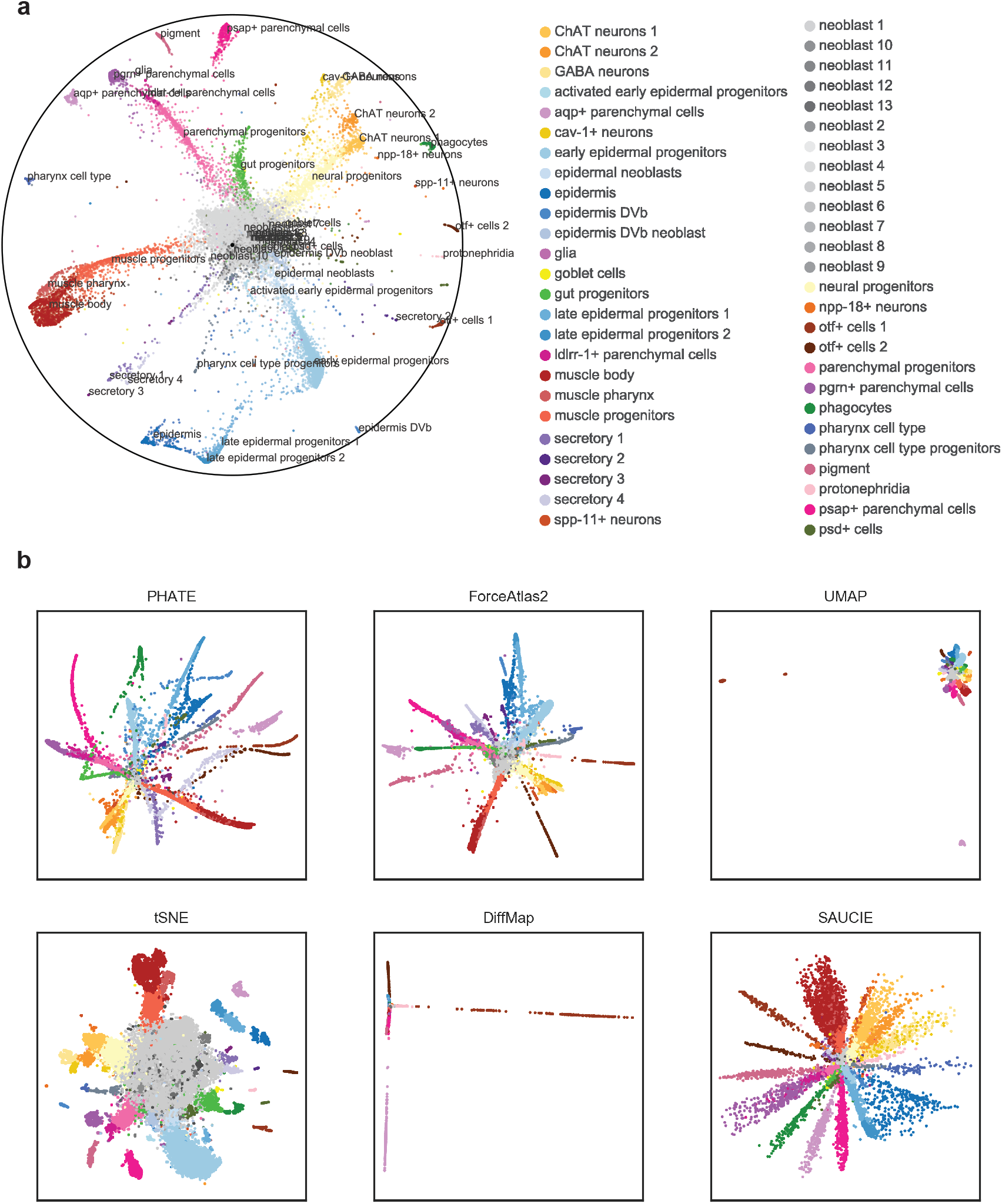
Comparison of various embeddings for the planaria Drop-seq dataset (Plass et al.). **(a)** Poincaré map rotated with respect to the root (medoids of neoblast 1 cluster). **(b)** Benchmark methods.

**Supplementary Figure 10.**
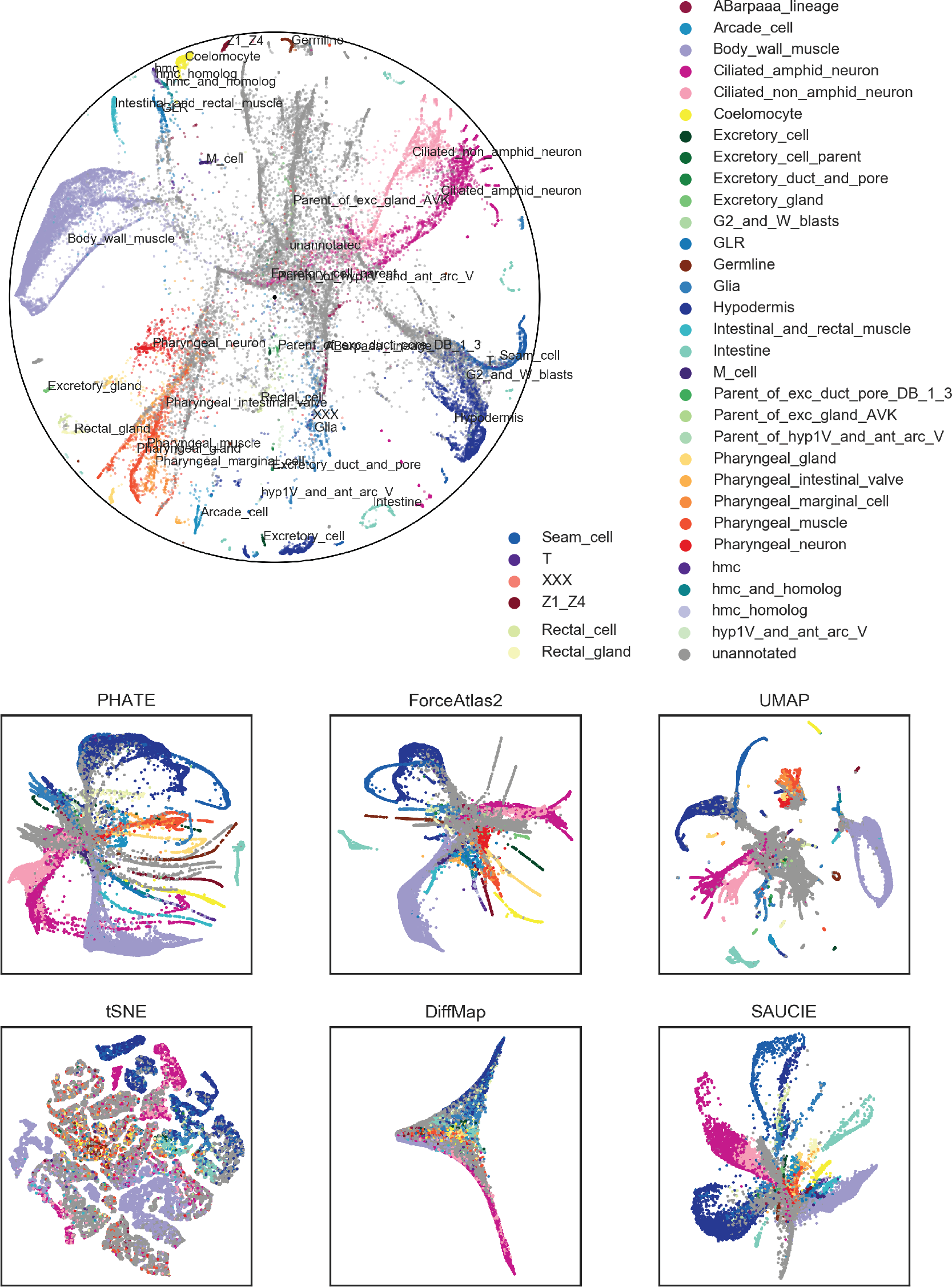
Comparison of various embeddings for the C. elegans 10X Genomics dataset (Packer et al.).

**Supplementary Figure 11.**
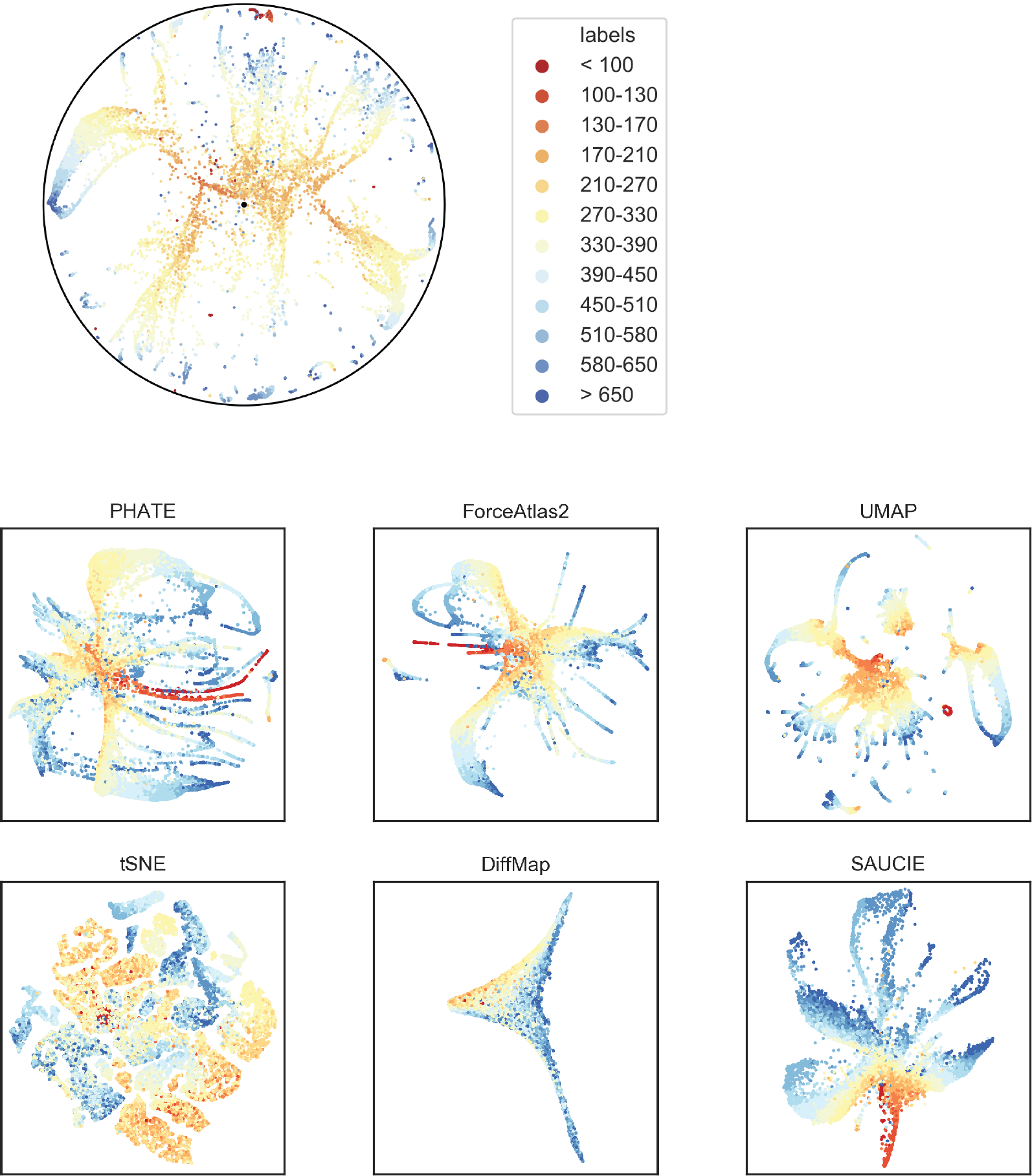
Comparison of the age of the embryo for various embeddings for the C. elegans 10X Genomics dataset (Packer et al.).

**Supplementary Figure 12.**
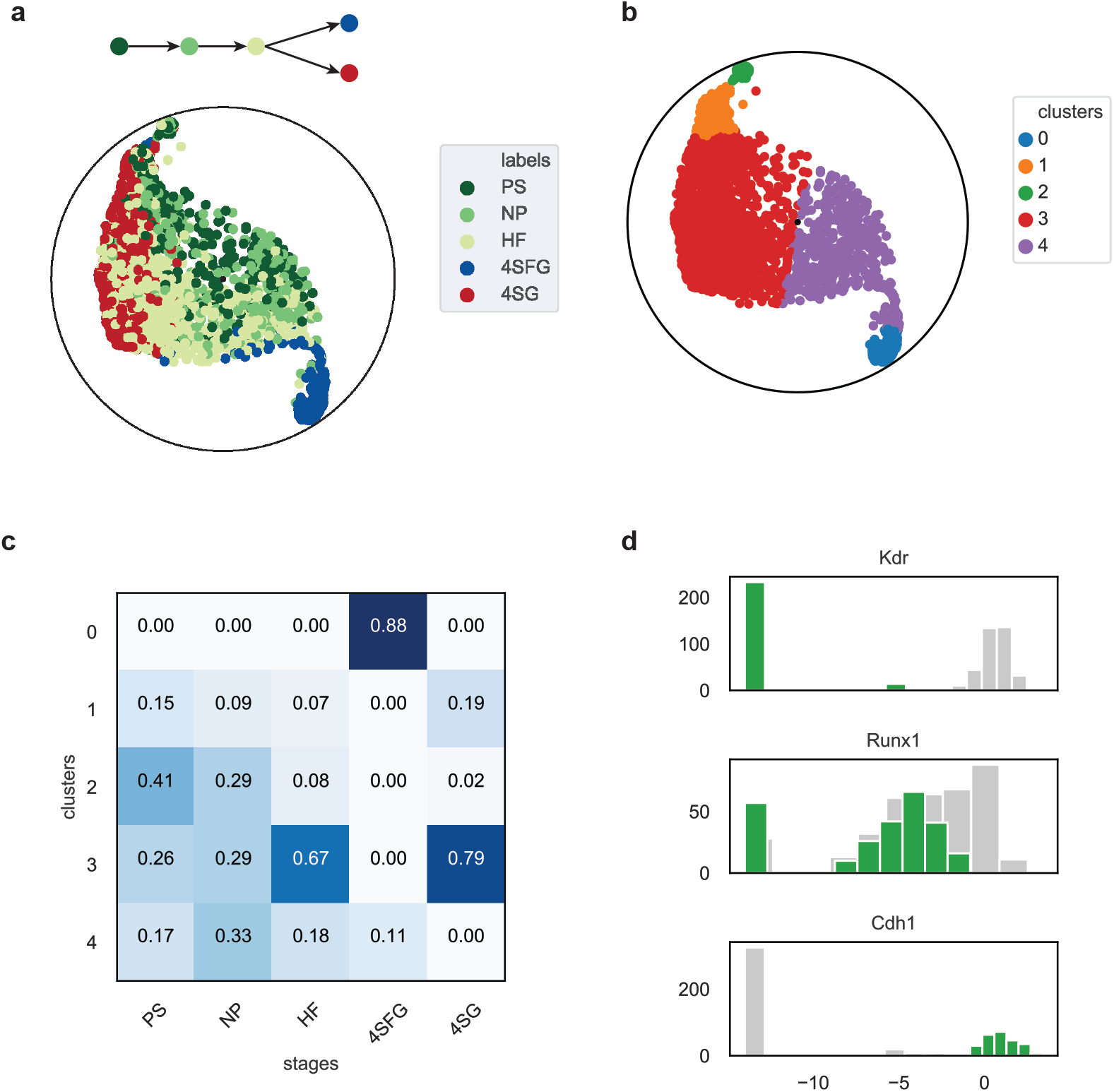
**(a)** Poincare map of the Moignard dataset. **(b)** Spectral clustering with Poincaré distances. **(c)** Analysis of stage-composition of the defined clusters. Clusters 1 and 3 most likely represent development of blood cells. Clusters 0 and 4 potentially correspond to endothelial development. Cluster 2 corresponds to the cluster named “mesodermal cells at primitive strike” in the original paper. **(d)** Comparison of the median expression of markers at PS stage for cluster 2 against the rest of PS cells. Cluster 4 consists mostly of Flk1-Runx1-.

**Supplementary Figure 13.**
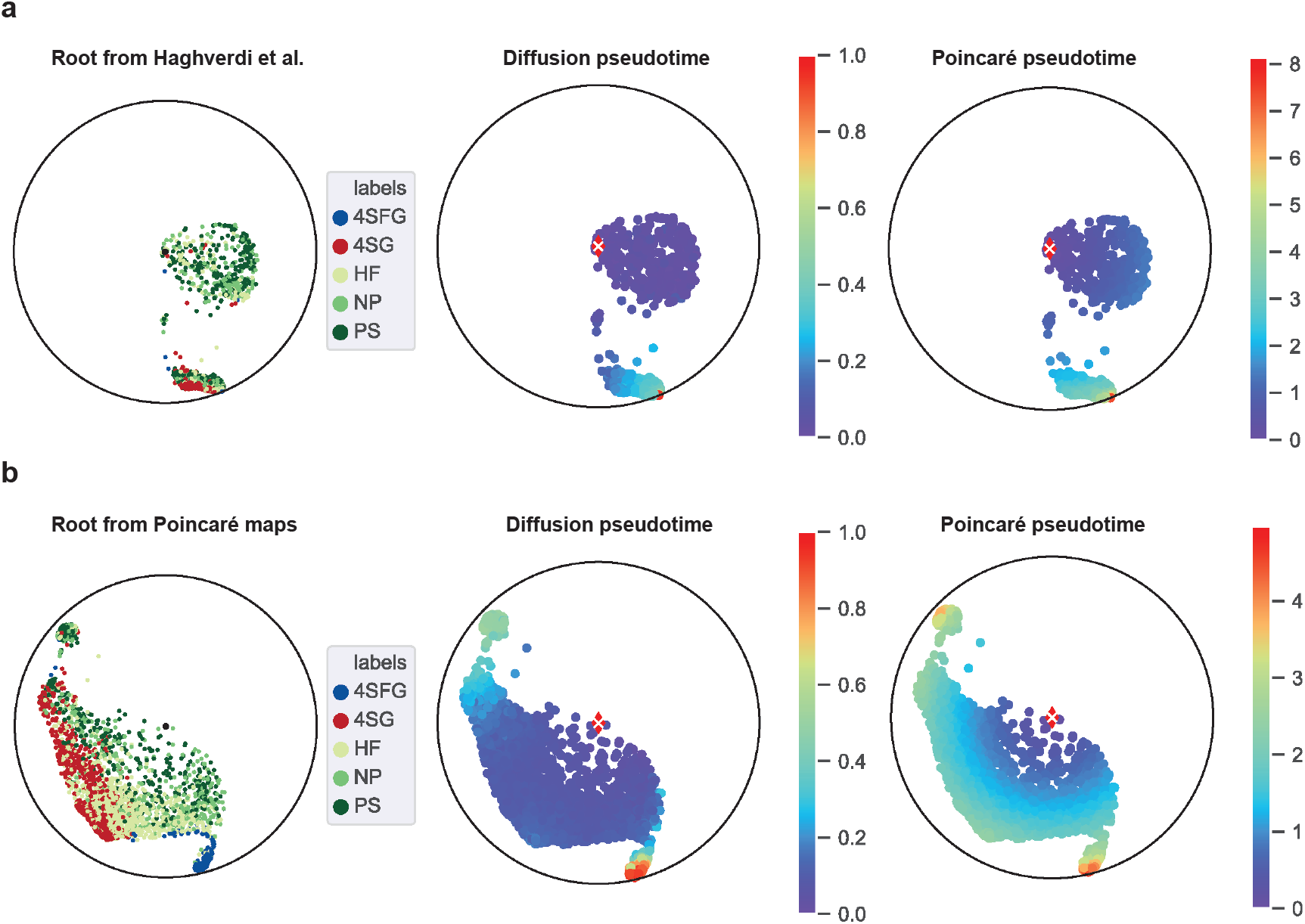
Rotation of the Poincaré map, and corresponding pseudotimes with root chosen according to **(a)** Haghverdi et al. **(b)** proposed new root.

**Supplementary Figure 14.**
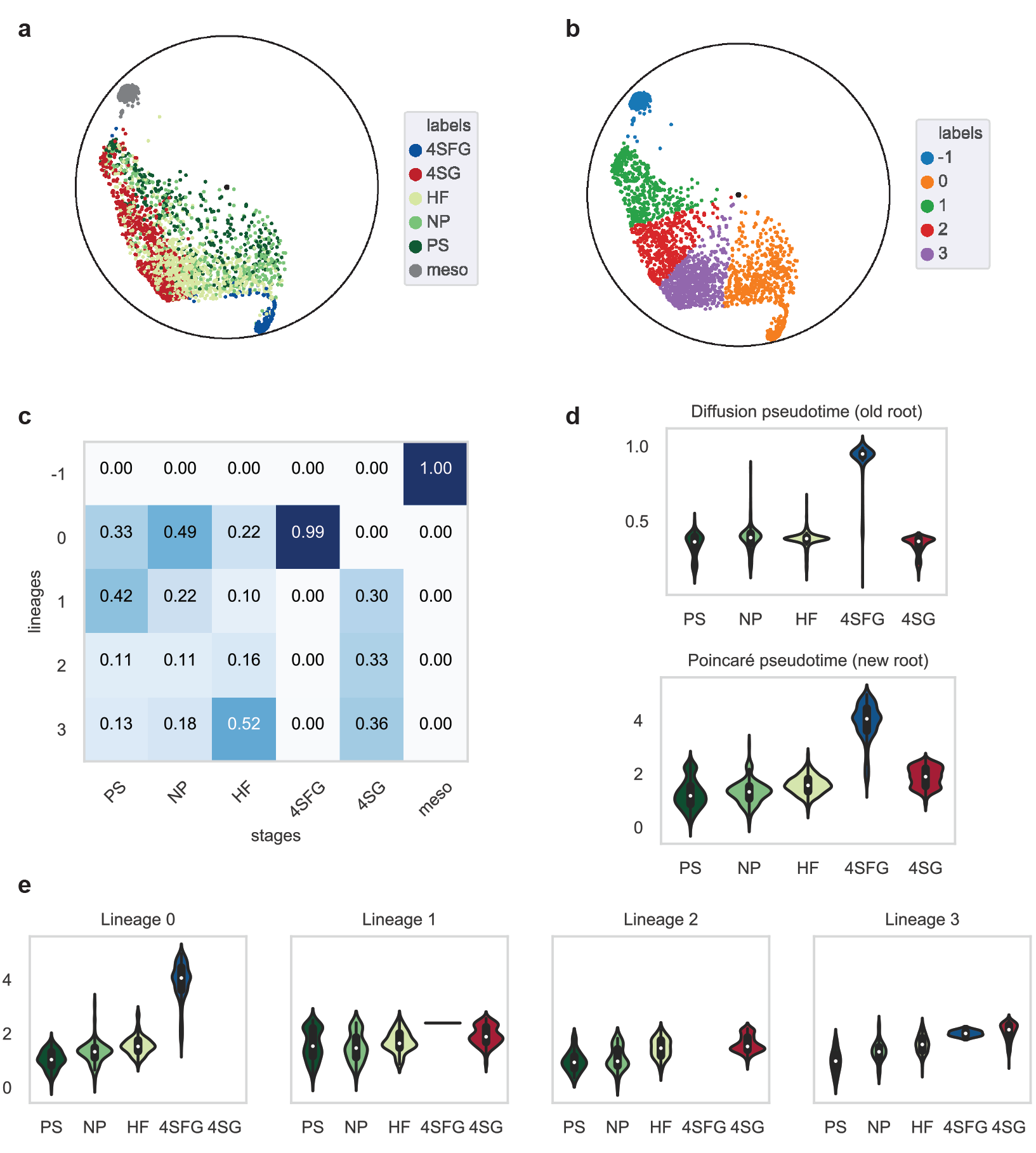
Analysis of stage ordering in different lineages. **(a)** Poincaré map with developmental stages. **(b)** Detected lineages with clustering by angle in the Poincaré disk. **(c)** Lineage composition per stage. **(d)** Average diffusion (from Haghverdi et al.) and Poincaré pseudotime per stage for the whole dataset. **(e)** Average Poincaré pseudotime per stage for in the individual lineage.

**Supplementary Figure 15.**
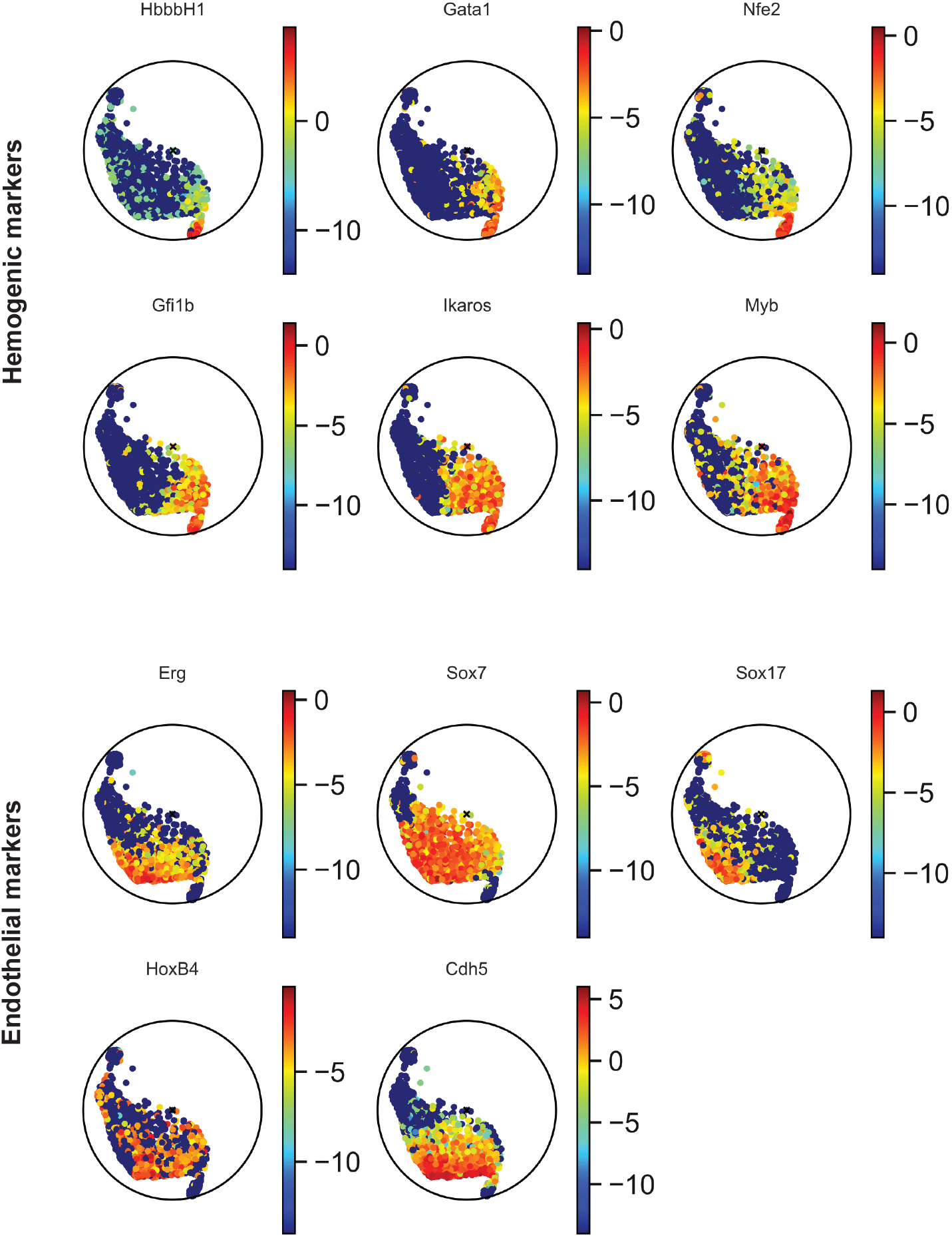
Expression of main endothelial and hemogenic markers visualized on the Poincaré disk.

**Supplementary Table 1.** Comparison of various clustering approaches on the synthetic datasets. Higher values mean better result.

| Dataset | ToggleSwitch |  | Myeloid progenitors |  | MP with blobs |  |
| --- | --- | --- | --- | --- | --- | --- |
| name | ARS | FMS | ARS | FMS | ARS | FMS |
| louvain | 0.46 | 0.59 | 0.58 | 0.63 | 0.89 | 0.91 |
| spectral Poincaré | 0.39 | 0.54 | 0.63 | 0.67 | 0.89 | 0.91 |
| agglomerative Poincaré | 0.49 | 0.61 | 0.59 | 0.64 | 0.89 | 0.91 |
| kmedoids Poincaré | 0.38 | 0.53 | 0.52 | 0.59 | 0.54 | 0.61 |
| spectral raw | 0.18 | 0.39 | 0.52 | 0.58 | 0.28 | 0.46 |
| agglomerative raw | 0.12 | 0.42 | 0.54 | 0.60 | 0.46 | 0.64 |
| kmedoids raw | 0.19 | 0.41 | 0.55 | 0.61 | 0.17 | 0.36 |
| spectral PCA | 0.18 | 0.39 | 0.53 | 0.59 | 0.29 | 0.41 |
| agglomerative PCA | 0.12 | 0.42 | 0.48 | 0.55 | 0.43 | 0.60 |
| kmedoids PCA | 0.20 | 0.42 | 0.49 | 0.56 | 0.65 | 0.70 |
| spectral tSNE | 0.47 | 0.60 | 0.59 | 0.64 | 0.85 | 0.88 |
| agglomerative tSNE | 0.36 | 0.51 | 0.49 | 0.56 | 0.89 | 0.90 |
| kmedoids tSNE | 0.43 | 0.57 | 0.43 | 0.51 | 0.71 | 0.76 |
| spectral UMAP | 0.37 | 0.52 | 0.42 | 0.50 | 0.89 | 0.91 |
| agglomerative UMAP | 0.37 | 0.52 | 0.52 | 0.58 | 0.89 | 0.91 |
| kmedoids UMAP | 0.31 | 0.48 | 0.58 | 0.63 | 0.66 | 0.71 |
| spectral DiffusionMaps | 0.55 | 0.66 | 0.52 | 0.58 | 0.76 | 0.81 |
| agglomerative DiffusionMaps | 0.04 | 0.39 | 0.11 | 0.31 | 0.12 | 0.44 |
| kmedoids DiffusionMaps | 0.06 | 0.36 | 0.42 | 0.50 | 0.33 | 0.50 |
| spectral ForceAtlas2 | 0.00 | 0.53 | -0.00 | 0.30 | 0.00 | 0.39 |
| agglomerative ForceAtlas2 | 0.48 | 0.61 | 0.56 | 0.61 | 0.89 | 0.91 |
| kmedoids ForceAtlas2 | 0.42 | 0.56 | 0.55 | 0.61 | 0.53 | 0.60 |

**Supplementary Table 2.** Comparison of diffusion pseudotime (dpt) and Poincaré pseudtotime (pmpt) against real time on synthetic datasets using Pearson correlation coefficient between. The last column corresponds to the correlation coefficient between diffusion pseudotime and Poincaré pseudtotime. Higher values are better.

| Dataset | dpt | pmpt | dpt-pmpt |
| --- | --- | --- | --- |
| ToggleSwitch: branch1 | 0.99 | 0.99 | 0.99 |
| ToggleSwitch: branch2 | 0.98 | 0.98 | 0.99 |
| ToggleSwitch: avg | 0.99 | 0.98 | 0.99 |
| MyeloidProgenitors: erythrocyt | 0.89 | 0.94 | 0.91 |
| MyeloidProgenitors: megakaryoc | 0.94 | 0.95 | 0.93 |
| MyeloidProgenitors: monocyte | 0.93 | 0.89 | 0.98 |
| MyeloidProgenitors: neutrophil | 0.91 | 0.91 | 0.99 |
| MyeloidProgenitors: avg | 0.92 | 0.92 | 0.95 |
| MyeloidProgenitors with blobs: Ery | 0.89 | 0.94 | 0.90 |
| MyeloidProgenitors with blobs: Mk | 0.94 | 0.96 | 0.93 |
| MyeloidProgenitors with blobs: Mo | 0.93 | 0.89 | 0.97 |
| MyeloidProgenitors with blobs: Neu | 0.91 | 0.91 | 0.99 |
| MyeloidProgenitors with blobs: avg | 0.92 | 0.92 | 0.95 |

## Footnotes

1 In general *k*NNG for a given *k* is not necessary connected. We need to enforce connectivity to reconstruct the hierarchy. If one component were disconnected from other components, it would be impossible to reconstruct its position relatively to other components.

